# Phosphoproteomics unveils the signaling dynamics in neuronal cells stimulated with insulin and insulin-like growth factors

**DOI:** 10.1101/2025.06.25.661584

**Authors:** Mereena George Ushakumary, Ryan Sontag, Camilo Posso, Thomas L Fillmore, Maisie E Topping, Heather M Olson, Philip L. De Jager, David A. Bennett, Zoe Arvanitakis, Vladislav A Petyuk

**Affiliations:** Bioq3logical Sciences Division, Pacific Northwest National Laboratory; Center for Translational & Computational Neuroimmunology, Department of Neurology and the Taub Institute for Research on Alzheimer’s Disease and the Aging Brain, Columbia University Irving Medical Center, New York City, NY, 10032, USA; Rush Alzheimer’s Disease Center, Rush University Medical Center, Chicago, IL, USA

**Keywords:** proteomics, phosphoproteomics, neural, insulin, IGF, Rho GTPase

## Abstract

**Background:** Given the role of metabolism in brain health and disease, investigating the role of insulin (INS) and insulin-like growth factors (IGFs) as potential therapeutic strategies for neurodegenerative diseases is currently underway. Yet, the signaling pathways associated with INS and IGFs in the brain remain elusive, particularly for the human brain. Unraveling these pathways is critical for harnessing their therapeutic potential in metabolism-associated brain disorders.

**Methods:** This study employed phosphoproteomics using human neuroblastoma cell line, SH-SY5Y, to unravel the signaling network of INS, IGF-1, and IGF-2. Briefly, cells were stimulated at 10 and 60-minutes with the ligands, followed by protein extraction, trypsin digestion, tandem mass tag (TMT)-labelling and phosphopeptides enrichment using an immobilized metal affinity chromatography (IMAC) and liquid chromatography-tandem mass spectrometry (LC-MS/MS) analysis. Data was processed using R statistical software. Protein annotations were obtained from the UniprotKB database, and pathway enrichment analysis was performed using Ingenuity Pathway Analysis (IPA).

**Results:** Phosphoproteomics performed at 10 and 60 minutes identified 34358 phosphosites of which 3284 were significant at 10 min and 2374 at 60 min (p.adj <0.05) across all three ligands. Ligand stimulation induced modulation in phosphorylation at both the receptor level and downstream signaling targets at serine (S), threonine (T) and tyrosine (Y) residues. Phosphorylation of LIMA1-Y229, a regulator of actin-cytoskeletal function, was the most prominent Y phosphosite across all ligands. IPA identified Rho GTPase, the molecular switches that regulate actin cytoskeletal dynamics, as the most significantly enriched pathway, with IGF-1 predominantly driving phosphorylation of Rho GTPase effectors such as Rho Guanine nucleotide exchange factors (ARHGEFs), Rho GTPase activating proteins (ARHGAPs) and CDC42. Myocardin related transcription factor A (MRTFA), a transcriptional target of Rho GTPase, was increased in ligand-stimulated cells at 10 min, and inhibition of Rho/SRF pathway by CCG1423 prevents nuclear localization of IGF-1-induced MRTFA.

**Conclusions:** This study demonstrates that INS, IGF-1 and IGF-2 regulate Rho GTPase and MRTFA activation, thereby contributing to the control of actin cytoskeletal dynamics in neuronal cells. Given the role of INS and IGFs in neuronal survival and neurodegenerative conditions, elucidating the mechanisms is of critical importance, as it offers insights into disease pathogenesis and potential therapeutic targets.

## Introduction

Phosphoproteomics is an advanced high-throughput technique designed to systematically profile protein phosphorylation, a pivotal regulatory mechanism in cellular signaling. Protein phosphorylation, mediated by the opposing actions of kinases and phosphatases, serves as a molecular switch to modulate protein activity, localization, stability, and interaction networks. These modifications are integral to intracellular communication, playing a central role in orchestrating numerous biological processes. The insulin/IGF signaling (IIS) pathway activates a cascade of phosphorylation events that regulate cellular growth and metabolism. In the context of IIS, phosphoproteomics emerges as a valuable tool for unraveling the complex web of phosphorylation events that underlie the critical signaling networks. Insulin and IGF-1/2 are peptide hormones that operate within intricate frameworks of ligands, receptors, and downstream pathways. Upon binding to their respective receptor tyrosine kinases-insulin receptor (IR) and IGF-1 receptor (IGF-1R)-these hormones trigger downstream signaling cascades, including the phosphoinositide 3-kinase (PI3K)-Akt and mitogen-activated protein kinase (MAPK) pathways^1–3^. These cascades regulate a wide array of cellular processes, such as glucose metabolism, protein synthesis, cell growth, and survival^4–6^. Effective regulation of these pathways is vital for the maintenance of cellular homeostasis.

In the brain, insulin signaling plays a central role in supporting neuronal function and metabolic processes by regulating glucose uptake and utilization^7–11^. This regulation is essential for maintaining neuronal energy homeostasis, which supports processes such as neurotransmission, synaptic plasticity, and overall brain function. IGF-1, in contrast, is primarily involved in neurodevelopment, where it promotes neuronal differentiation, axonal outgrowth, and myelination^12^. Recent research has also underscored the significance of IGF-2 as a key regulator of learning, memory, and synaptic plasticity^13^. Together, insulin and IGF signaling pathways are crucial for neuronal health, supporting cognitive functions such as learning and memory while providing protection against cellular stress and damage. Resistance to insulin and IGFs, a well-known impairment of the pathway, can occur under a variety of conditions, including obesity, physical inactivity, genetic predisposition, certain medications, hormonal imbalances, and aging, often with overlapping contributing factors^14–21^. Disruption in these essential signaling events, affecting their ability to activate downstream cascades such as the PI3K-Akt and MAPK pathways leads to glucose metabolism defects, oxidative stress, and inflammation-factors commonly associated with metabolic disorders like type 2 diabetes and neurodegenerative diseases such as Alzheimer’s disease (AD) and cerebrovascular dementia^9,22,23^. Numerous conditions associated with neurodegeneration and cognitive decline, such as insulin resistance, amyloid-beta buildup, and tau hyperphosphorylation, share common signal transduction abnormalities at the molecular level^24^. Understanding the molecular mechanisms governing these pathways can provide potential therapeutic targets, such as enhancing insulin sensitivity or modulating IGF-1 activity, to mitigate brain-related diseases and promote cognitive health. Phosphoproteomics has been used to distinguish signaling differences between IR and IGF-1R in preadipocytes, revealing that IR preferentially activates mTORC1 and Akt pathways, while IGF-1R favors Rho GTPase and cell cycle signaling^3^. In neuronal cells, phosphoproteomics has enabled temporal mapping of synaptic signaling and stress responses, including studies in cortical and hippocampal neurons, and neuroblastoma cell lines (SH-SY5Y, IMR-32) under various stimuli such as retinoic acid and oxidative stress^25–28^. However, a comprehensive temporal phosphoproteomics analysis of neuronal cells in response to INS, IGF-1, and IGF-2 is currently lacking; an important gap in understanding the fundamental signaling dynamics these peptides elicit in neurons.

In this study, we aimed to investigate the molecular dynamics of INS, IGF-1, and IGF-2 signaling in human SH-SY5Y neuroblastoma cells^29^, which exhibit neuronal-like morphology and express human-specific proteins and protein isoforms. To achieve this, we employed LC-MS/MS–based phosphoproteomics analysis on SH-SY5Y cells stimulated with INS, IGF-1, and IGF-2 at 10 and 60 minutes and analysed the ligand-specific and temporal phosphorylation patterns in these cells. We then conducted pathway enrichment analysis to elucidate the signaling networks and biological processes affected by ligand-specific and temporal phosphorylation changes.

## Methods

### Cell culture

SH-SY5Y neuroblastoma cells were cultured using the standard ATCC-recommended EMEM medium containing 10% FBS. The cells were serum-deprived prior to stimulation with recombinant human INS (Cat# 12585-014), recombinant human IGF-1 (Cat# PHG0071), and recombinant human IGF-2 (Cat# PHG0084). Cell lysates were collected at various time points during preliminary studies, and 10 minutes and 60 minutes were selected for phosphoproteomic analysis based on the observed early activation and subsequent decline of phospho-Akt with INS and IGF-2 (Supplementary Fig 1). Nuclear extraction of IGF-1-stimulated and non-stimulated SH-SY5Y cells was carried out using NE-PER Nuclear and Cytoplasmic Extraction Reagents (Cat# 78833) following the manufacturer’s protocol. The chemical inhibitors used in this study included Rhosin hydrochloride (Cat# 5003), CCG 1423 (Cat# 5233), and Wortmannin (Cat# 1232). These inhibitors were added to the cell culture medium 30 minutes prior to stimulating the cells with IGF-1, at concentrations of 50 µM, 10 µM, and 50 nM, respectively.

### Western Blotting

SH-SY5Y cells treated with or without INS, IGF1 and IGF2 were harvested, and protein abundance was determined by western blot analysis following standard protocols. Briefly, the lysates were electrophoresed and then transferred onto a PVDF membrane. The membranes were blocked and incubated overnight at 4^º^C with the following primary antibodies: anti-phospho AKT473 and AKT, MRTFA (E3N7W, 77098S), and PCNA (10205-2-AP). Enhanced chemiluminescent reagent was used to detect the proteins and imaged the blots using Bio-Rad ChemiDoc Imaging System. The target proteins were normalized with beta actin for cytoplasmic proteins and Proliferating cell nuclear antigen (PCNA) for nuclear proteins. Target protein expression was quantified by densitometric analysis of scanned blots using ImageJ software following Bio-Rad’s image analysis and quantification method^30^. Briefly, western blot bands were selected using the rectangular selection tool and defined each band by numbering them. A histogram was generated for each band and a straight-line selection tool was used to draw the line on the bottom of picks to make it close completely. Wand tool from ImageJ generated the area and percent values for the histograms and generated the excel sheets, which was used to analyze the fold change of protein of interest compared to the internal controls. Fold change was calculated over control samples.

### Phosphoproteomics sample preparation

#### Protein Digestion and Cleanup

SH-SY5Y cell pellets were digested as previously described^31,32^. Briefly, samples were resuspended in 650 uL of fresh lysis buffer containing 4% (wt/vol) sodium deoxycholate (SDC) and 100 mM Tris-HCl, pH 8.5, heated at 95°C for 5 minutes, and lysed by probe sonication. Reduction/alkylation buffer containing 100 mM tris(2-carboxyethyl) phosphine (TCEP) and 400 mM 2-chloroacetamide (CAM) (pH 7) was added at 1:9 ratio (vol/vol) and incubated at 45°C for 10 minutes. Samples were digested overnight with Lys-C (1 mAU per 50 ug protein) and trypsin (1:50 wt/wt) at 37°C. The digestion reaction was quenched and SDC precipitated by acidifying to 1% formic acid (FA). Undigested debris and the SDC were pelleted by centrifugation at 25,000 rpm for 10 minutes and the supernatant was desalted using Phenomenex 100 mg C18-E solid phase extraction (SPE) columns^33^. A BCA protein assay was used to measure peptide concentration.

#### TMT Labeling

An aliquot of 400 ug of peptides from each sample was dried down and resuspended in 80 uL of 50 mM HEPES (pH 8.5) for a 5 ug/uL concentration. TMT10 reagents (Thermo Fisher Scientific) were reconstituted to a 20 mg/mL concentration in anhydrous acetonitrile and added to the peptide aliquots at a 1:1 ratio (wt/wt). The reactions were incubated for 1 hour at 25°C in a thermomixer shaking at 400 rpm. A 3.2 ug aliquot was taken from each sample in the plex, combined, and diluted to 100 uL with 0.1% FA. This premix QC was cleaned with C18 stage tips^33^ and submitted for LC-MS/MS analysis to determine the success of the labeling reactions. The remaining samples were frozen until this analysis was complete. Once the TMT labeling success was confirmed, the samples were diluted to 2.5 ug/uL with 50 mM HEPES pH 8.5, 20% ACN and the reactions were quenched by adding 12 uL of 5% hydroxylamine and incubating at 25°C for 15 minutes with 400 rpm shaking. After quenching, the samples in each plex were combined and dried down completely, then reconstituted in 1 mL of 3% ACN, 0.1% FA and desalted using a 200 mg C18 column. The sample was fractionated using high pH reverse phase chromatography on an Agilent Zorbax column and concatenated into 12 fractions. Each fraction was acidified to 0.1% FA and 5% of the resulting sample was removed for global proteomics analysis.

#### Phosphopeptide Enrichment

Phosphopeptides were enriched by Fe3+ immobilized metal affinity chromatography (IMAC) as previously described^33^ with the following alterations. Following the 30-minute incubation, the beads were washed with 1 mL of 80% ACN, 0.1% trifluoroacetic acid (TFA) before being transferred to equilibrated C18 stage tips. For LC-MS analysis, the Phosphopeptides were reconstituted in 12 uL of 3% ACN, 0.1% FA.

#### LC-MS/MS Mass spectrometry

Proteomic analyses of 5 uL of reconstituted phosphopeptide solution were performed using a Dionex UltiMate system (Thermo Scientific) coupled to a Q-Exactive HF-X mass spectrometer (Thermo Scientific). Peptides were first passed through a trapping column (150 um inner diameter (i.d.) x 2 cm length, packed with 5 um pore size Jupiter C18) at 7 uL/min for 2 minutes with Mobile Phase A (0.1% formic acid (FA) in water). Peptides were eluted from the trapping column to the analytical column (Waters Acquity BeH 130, 1.7 um pore size, 75 um i.d. x 20 cm length) operated at a temperature of 45°C. Chromatographic separation was performed at 0.2 ul/min with the following gradient: 1–8% (0 min–7 min), 8–21% (7 min–82 min), 21–28% (82 min–102 min), 28–37% (102 min–112 min), 37–75% (112 min–117 min) and 75%–95% (117 min–120 min) Mobile Phase B (0.1% FA in acetonitrile). The ion transfer tube temperature was set to 300 °C, and the nano electrospray voltage was 2.2 kV. Each dataset was collected over 100 minutes, following a 10-minute delay after sample trapping and the start of the gradient. FT-MS spectra were acquired in the range of 300 to 1800 m/z at a resolution of 60 k (AGC target 3e6), while the top 12 FT-HCD-MS/MS spectra were acquired in data-dependent acquisition (DDA) mode with an isolation window of 0.7 m/z at a resolution of 45 k (AGC target 1e5). HCD fragmentation was performed with a normalized collision energy of 30, using a 45-second exclusion time and analyzing charge states 2 to 6.

### Data analysis

Tandem Mass Tag (TMT) labeling-based phosphoproteomics data were processed and analyzed using MSGF+ and MASIC^34,35^. The data were searched against the human reference proteome from UniProt^36^ with appropriate modifications for TMT labeling and phosphorylation. Quantitative analysis was performed using the reporter ion intensities for each TMT channel. We filtered the data such that we have at least 50% of the data present per phospho-site. We also performed batch correction using ComBat^37^, to moderate any batch effects. Differential phosphorylation analysis was conducted using the Limma package in R, and significance was determined based on a corrected p-value cutoff of <0.05. Data visualization with bubble heatmaps was performed with the *enrichmap* function from the ‘TMSig’ R package^38^. The Massive ID to access the raw dataset is MSV000097782.

### Ingenuity pathway analysis (IPA)

For phosphoproteins showing significant changes in phosphorylation levels in ligand-stimulated cells compared to controls, pathway and functional enrichment analyses were conducted using Ingenuity Pathway Analysis (IPA) to identify enriched biological processes and signaling pathways. Networks automatically generate plausible signaling cascades, describing potential mechanisms of action that lead to observed gene expression changes from the data. The dataset was formatted according to the input requirements of the IPA software. This involved organizing the data into tabular format, ensuring the inclusion of identifiers such as gene symbols, protein names, and corresponding phosphorylation sites with their relative abundances or fold changes. Appropriate settings were configured for the analysis, including species (Homo sapiens) and reference set (Ingenuity knowledge base). A core analysis was conducted in IPA to identify biological pathways, networks, and functional categories significantly affected by the phosphorylation changes. During the analysis, parameters such as p-value cut-off (p. adj< 0.05) was set to determine significance.

## Results

### Global phosphoproteome changes

INS, IGF-1 and IGF-2 stimulation of SH-SY5Y cells showed an increase in AKT phosphorylation 10 minutes after stimulation, which subsided over time (Supplementary Fig. 1). A concentration of 0.1 nM ligands was sufficient to elicit this response. For the phosphoproteomics study, we used 0.1 nM concentrations of ligands and stimulated the cells for 10 and 60 minutes, including corresponding unstimulated vehicle controls. At the global level, a total of 8844 proteins were identified (Supplementary dataset 1). Following the ligand stimulation, 34,358 phosphosites, were identified across INS, IGF-1, and IGF-2 at 10 and 60 minutes compared to controls (Supplementary dataset 1). A general reduction in phosphorylation was observed upon ligand stimulation at 60 minutes compared to 10 minutes.

### Ligand-specific and temporal dynamics of stimulation

SH-SY5Y cells exhibited distinct phosphorylation response to individual ligands over time (Supplementary dataset 1). Based on the data, 2,240 significant phosphosites were identified following insulin (INS) stimulation, 1,991 with IGF1 stimulation, and 1,524 with IGF2 stimulation at the 10-minute mark. Among these, 203 phosphosites overlapped across all three ligands at the early time point (Fig. 2A). By 60 minutes, the numbers shifted to 1,410 for INS, 2,429 for IGF1, and 1,146 for IGF2 stimulation with 586 overlapping sites across all three ligands (Fig. 2B). These findings suggest that INS and IGF2 primarily triggered immediate activation of phosphosites, whereas IGF1 displayed a more sustained effect, with more phosphorylation sites at 60 minutes. The overlapping sites identified included both significantly upregulated and downregulated phosphosites. Additionally, temporal dynamics also revealed overlapping phosphosites between the 10- and 60-minute time points within each ligand, with IGF1 exhibiting the highest overlap (998 phosphosites) compared to INS and IGF2 (Fig. 2C). These observations clearly demonstrate that phosphorylation dynamics depend on both the ligand itself, as well as the timing of stimulation in SH-SY5Y cells.

**Figure 1:**
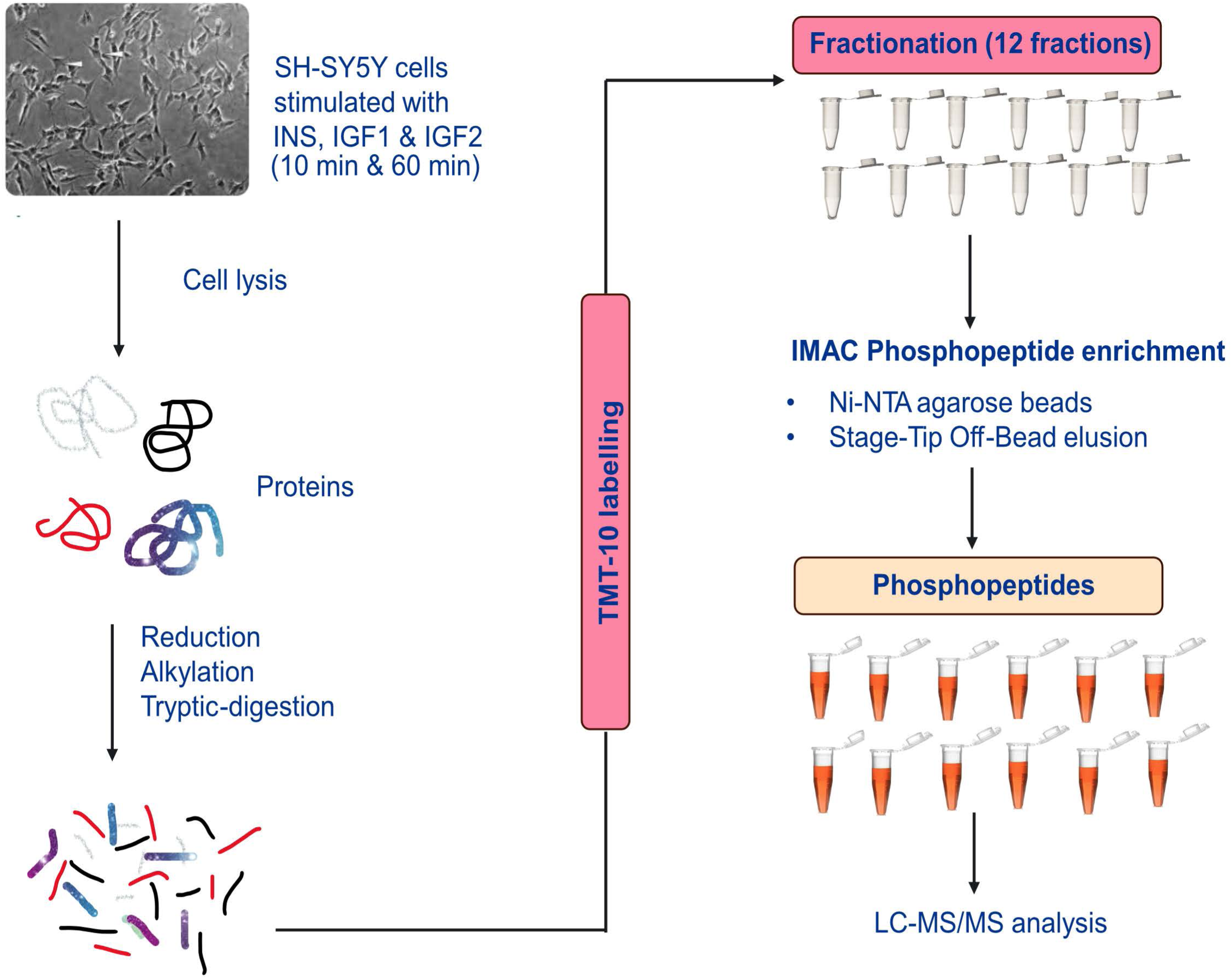
Summary of the experimental system and design. SH-SY5Y cells were stimulated with INS, IGF-1, & IGF-2, or unstimulated for 10- and 60-min. Cell lysates were collected, extracted proteins, and enzymatically digested them to peptides. They were fractionated into 12 fractions after labeling with Tandem mass tags (TMT). The Phosphopeptides were enriched using Ni-NTA agarose beads and analyzed using LC-MS/MS. Six replicates were used to perform individual ligand stimulation.

**Figure 2:**
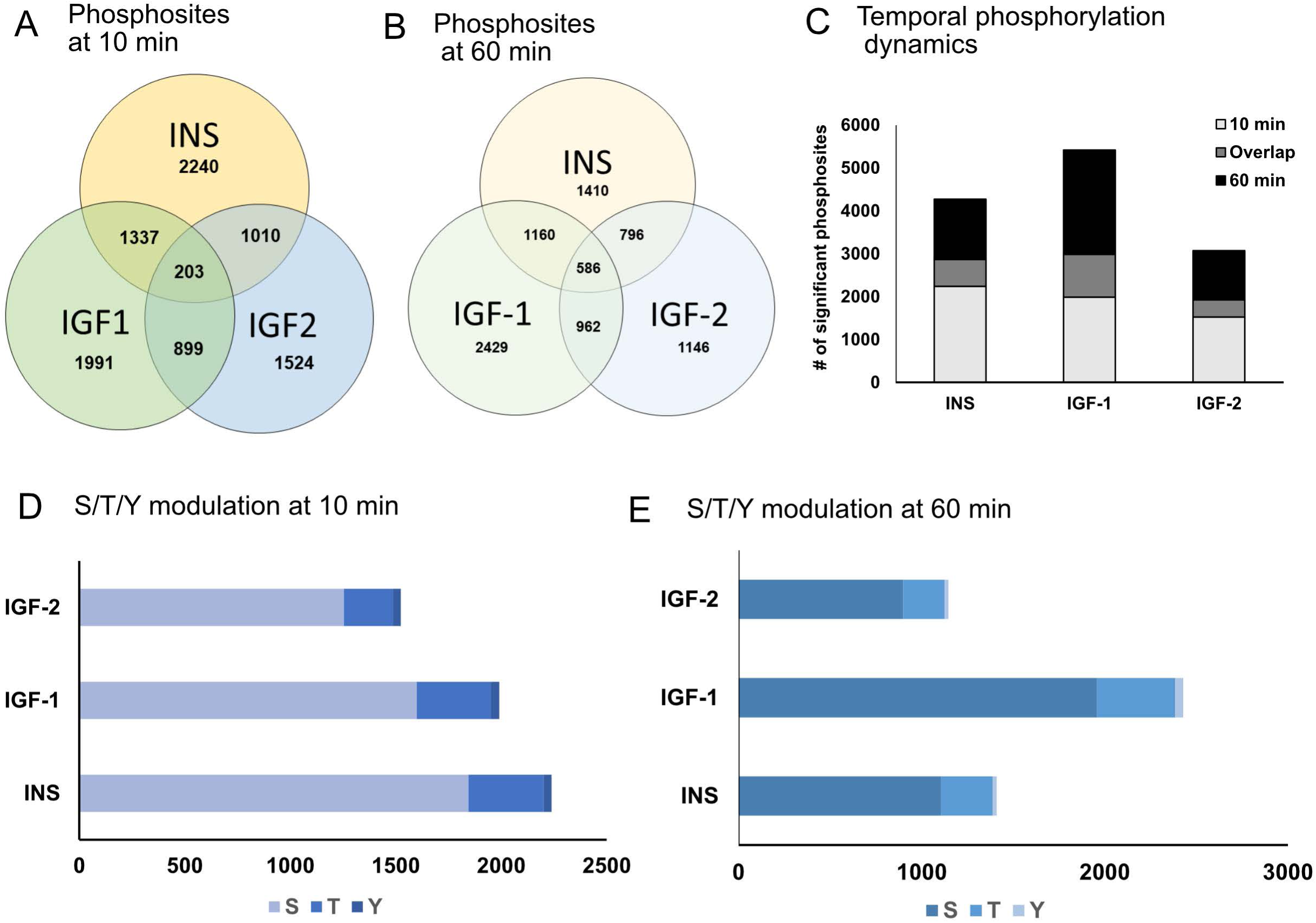
Global phosphoproteomics analysis. (A) Phosphorylation dynamics of INS, IGF-1 and IGF-1 at 10 min. The Ven diagram represents the individual changes in number of significant phosphosites within each group as well as the overlapping sites between the ligands. (B) Phosphorylation dynamics of INS, IGF-1 and IGF-1 at 60 min. Again, the Ven diagram illustrates the individual changes in the number of significant phosphosites within each group, as well as the overlapping sites between the ligands. IGF-1 shows a distinct stimulation pattern compared to INS and IGF-2. (C) Temporal dynamics in phosphorylation within each ligand at both time points. Temporal overlap of phosphosites were also more and unique with IGF-1. (D) Ser/Thr/Tyr phosphorylation dynamics at 10 min across all ligands. (E) Ser/Thr/Tyr phosphorylation dynamics at 60 min across all ligands. Ser changed from 82 to 80 % over time and Thr from 16 to 18%. However, Tyr (2%) remained the same across the time points.

### Dynamics of Ser/Thr/Tyr (S, T, Y) phosphorylation

Phosphorylation changes in S, T, and Y residues were plotted, revealing as expected, the highest number of significant phosphosites for S, followed by T and Y, regardless of ligand or time (Supplementary Figure 2). Ligand-specific changes indicated that INS induced more significant S and T phosphosites at 10 minutes compared to IGF-1 and IGF-2 (Fig. 2D). However, at 60 minutes, IGF-1 showed more significant S, T, and Y sites (Fig. 2E). We further analyzed proteins with significant phosphosites upon ligand stimulation at both time points, which were clustered together and visualized as heatmaps (Supplementary Figures 3 & 4). The top 20 significant phosphosites identified from INS, IGF-1, and IGF-2 stimulation at 10 and 60 minutes respectively are summarized (Fig. 3). PATL-T194, an mRNA decay factor, was consistently the top identified phosphosite in all three ligands at both time points. Additionally, RPS6KB1-S452, AT1S1-T246, RICTOR-T1135, PDCD4-S457, MTOR-T2474 LIMA1-S225, among others, were significantly higher at 10 min in all three ligands. At 60 min, IGF-1 showed a prominent increase in phosphosites such as CAD-S1859, STIP1-S16, ACIN1-S240, LAMTOR1-S98, ARFGEF2-S277, with slightly lower but essentially a similar response between INS and IGF-2. CAD-1859 and STIP1-S16 are phosphorylation targets of MTORC1.

**Figure 3:**
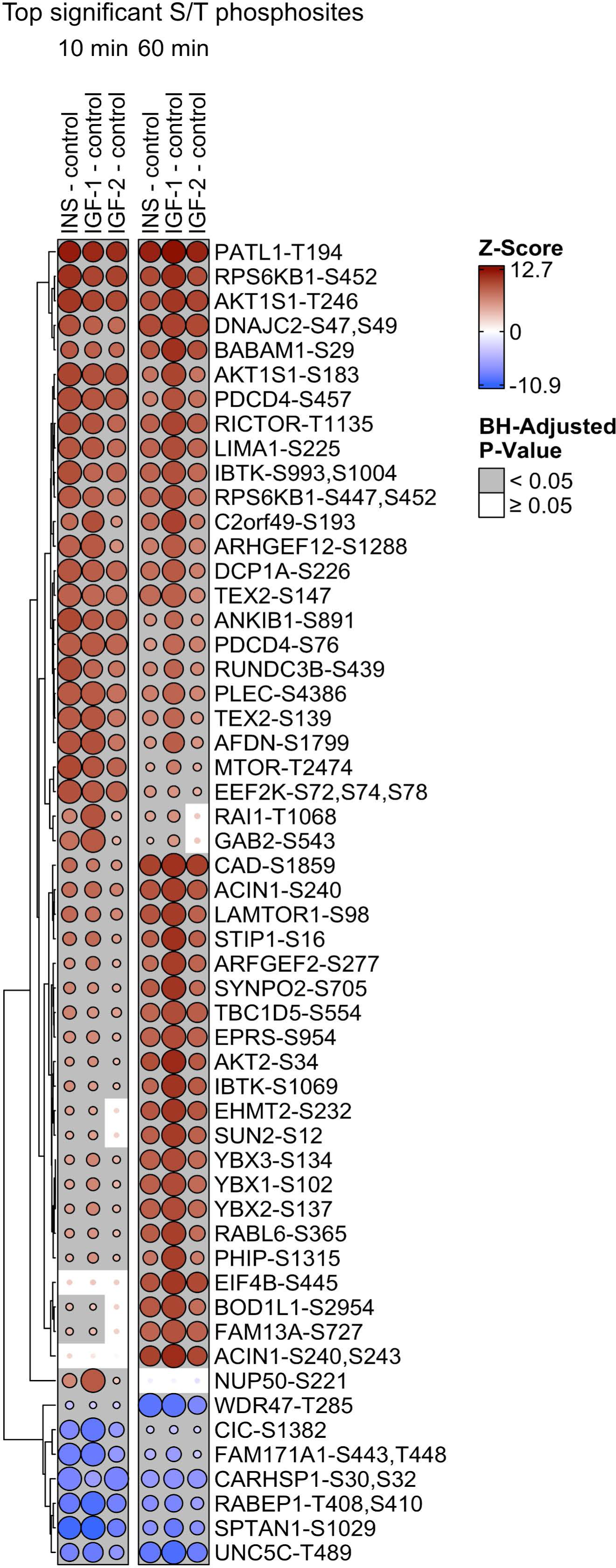
S/T dynamics upon ligand stimulation. The bubble heatmap represents the top significant S and T phosphosites upon stimulation with INS, IGF1 & IGF2 at 10- and 60-min. Bubbles are scaled by row relative to the most significant contrast. The bubble heatmap was created with the *enrichmap* function from the ‘TMSig’ R package.PATL-T194 was the topmost site identified and was significant higher in INS and IGF-stimulated cells at both time points. A prominent increase of CAD-S1859, ACIN-S240, LAMTOR-S98, STIP-S16 etc were found at 60 min with a prominent increase in IGF-1-treated cells.

### Y phosphorylation at the receptor level

In addition to S and T phosphosites, we also explored Y phosphosites at the receptor level. Among the total significant Y sites, phosphorylation at the receptor level was minimal. There was no significant phosphorylation sites observed for IGF-2 at the receptor level at either 10 or 60 minutes (Fig. 4A & B). INSR-Y1185 and IGF1R-Y1161 were found in both INS and IGF-1 stimulated cells at 10 minutes (Fig. 4A), and only IGF-1 maintained phosphorylation of these Y residues at 60 minutes (Fig. 4B). In addition to these canonical receptors, we also identified Y phosphorylation at position 1507 in anaplastic lymphoma kinase (ALK). ALK is known to be activated in cancer, and given the SH-SY5Y cell line nature, it’s challenging to directly correlate this phosphorylation to the ligands.

**Figure 4:**
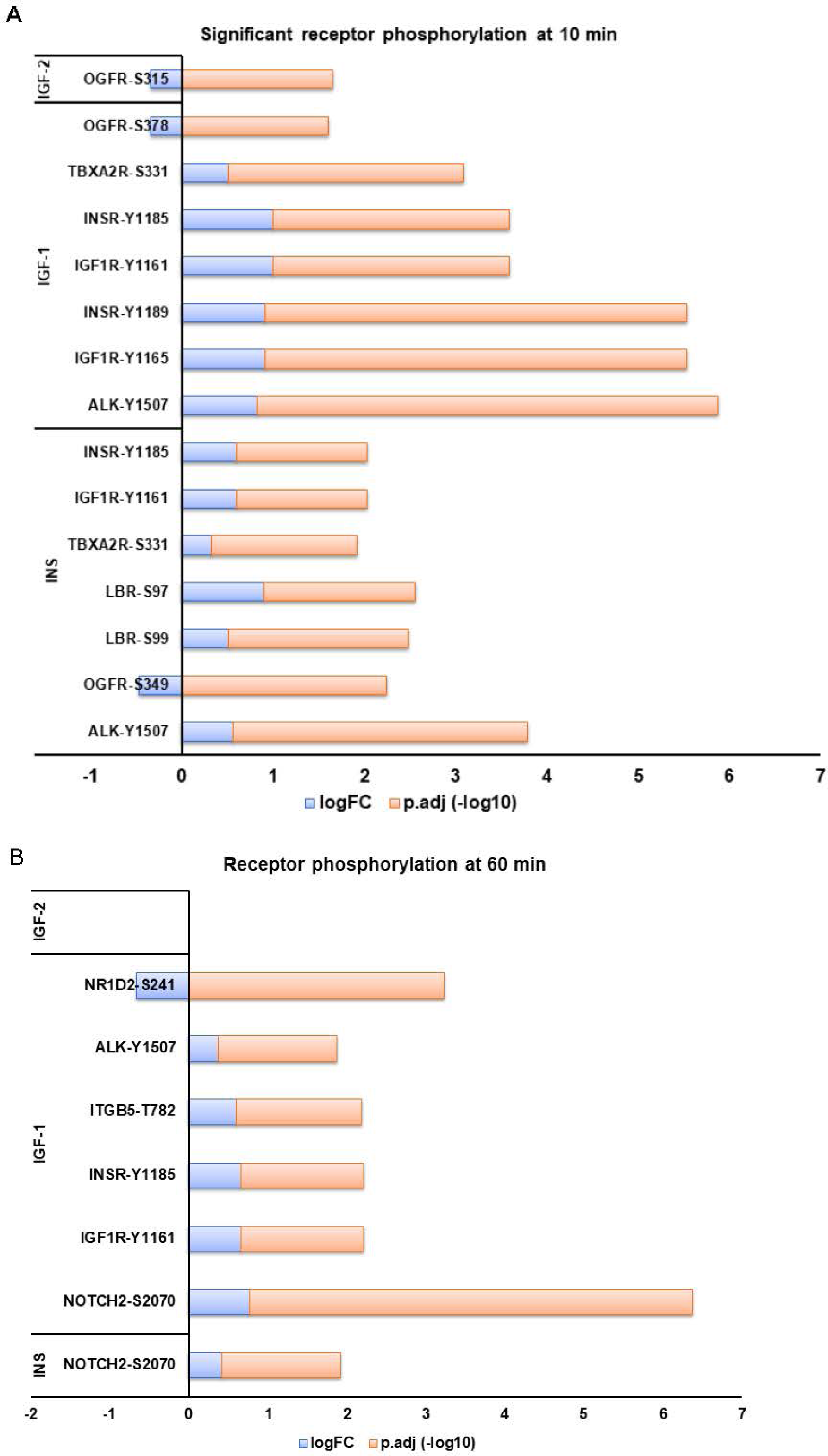
Tyrosine phosphorylation of receptors at 10 and 60 min. (A) Significant receptor tyrosine sites were extracted from the 10 min time point and plotted here for INS, IGF-1 and IGF-2. Different colors at X axis define the LogFC and -log10 adjusted P value. (B) Significant receptor tyrosine sites were extracted from the 60 min time point and plotted here for all the ligands. Different colors at X axis define the LogFC and -log10 adjusted P value. None of the receptors showed significant tyrosine phosphorylation upon IGF-2 stimulation at 60 min.

### Y phosphorylation at the downstream molecules

At the downstream level, LIM domain and actin-binding protein 1 (LIMA1)-Y229, involved in actin cytoskeleton regulation and dynamics, was the topmost Y phosphosite identified at 10 and 60 min across all three ligands (Fig. 5A). To elucidate the role of LIMA1-Y229, the most significant Y phosphosite, in INS and IGF stimulation, we utilized STRING database^39^ to identify its functional interaction partners (Supplementary figure 5A). Based on the STRING database, the predicted functional partners of LIMA1 include FLNA, FLNB, RHOA, VCL, TJP, CTNNA1, CTNNB1, CTNND1, CDH1 and CDH17. These genes correspond to actin binding proteins involved in actin cytoskeleton and dynamics^40,41^. We identified significantly modulated phosphosites associated with these genes upon ligand stimulation and these were visualized using heatmaps for 10 min and 60 min collected data (Supplementary figure 5C & D). Both time points show an increase in phosphorylation of the LIMA-interacting proteins in INS, IGF-1 and IGF-2 groups, compared to controls. At 10 min, Y113 site of Pre-mRNA 3’-end-processing factor FIP1 (FIP1L1-Y113) emerged as another significant site with INS and IGF-1 showing the stronger response. Phosphorylation of FIP1L1 at the Y113 position has not been previously reported, making this the first instance of INS and IGFs inducing Y113 phosphorylation on FIP1L1 in SH-SY5Y cells. The canonical MAPK phosphorylation (MAPK3 & MAPK1) was significant only in INS and IGF-1 groups at 10 min with a stronger response in IGF-1 stimulated neuronal cells. Moreover, the phosphosites MAPK1-Y187, IGF1R-Y1165 and INSR-Y1189 were only significant in IGF-1 stimulated cells at 10 min making it very specific to IGF-1 mediated events.

**Figure 5:**
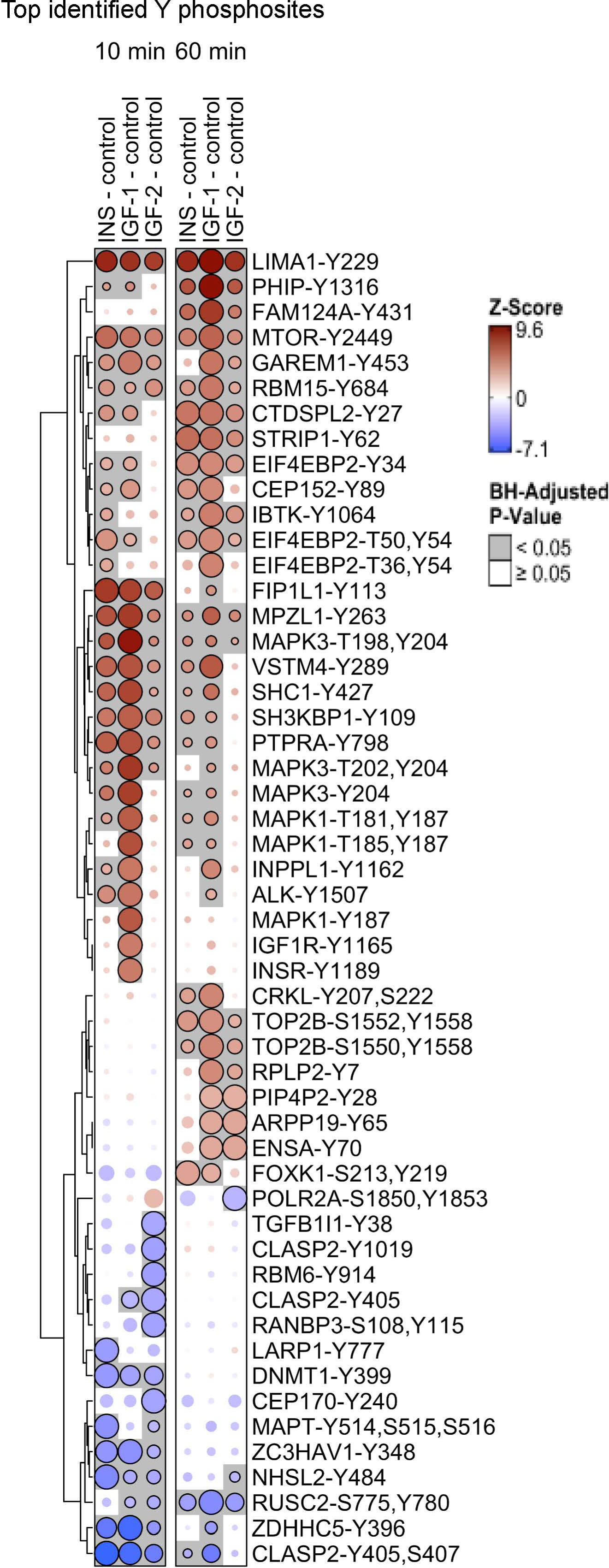
Tyrosine phosphorylation at the downstream targets. (A) The bubble heatmap represents the top significant Y phosphosites identified upon stimulation with INS, IGF-1 & IGF-2 at 10- and 60-min. Bubbles are scaled by row relative to the most significant contrast. The bubble heatmap was created with the *enrichmap* function from the ‘TMSig’ R package. LIMA1-Y229 was the topmost phosphosite identified across all the ligands at both time points. At 10 minutes, FIP1L1-Y113 exhibited a strong response across all ligands. Notably, MAPK phosphosites identified at this time point were predominantly upregulated by IGF-1.

### Ingenuity pathway analysis

To understand the biological insights from the phosphorylation data, we conducted a pathway enrichment analysis for the identified genes and phosphosites at 10 and 60 min. For INS, the top significant pathways identified were the Rho GTPase cycle, IR signaling, Processing of Capped intron-containing pre-mRNA, HER-2 signaling, mTOR signaling, and Reelin signaling (Fig. 6A). Except for the Processing of Capped intron-containing pre-mRNA and HER-2 signaling, the rest remained the same for IGF-1, with the addition of FLT3 and NGF signaling. IGF-2 also showed enrichment in Rho GTPase cycle, IR signaling, and mTOR signaling along with HER-2 and 14-3-3-mediated signaling.

**Figure 6:**
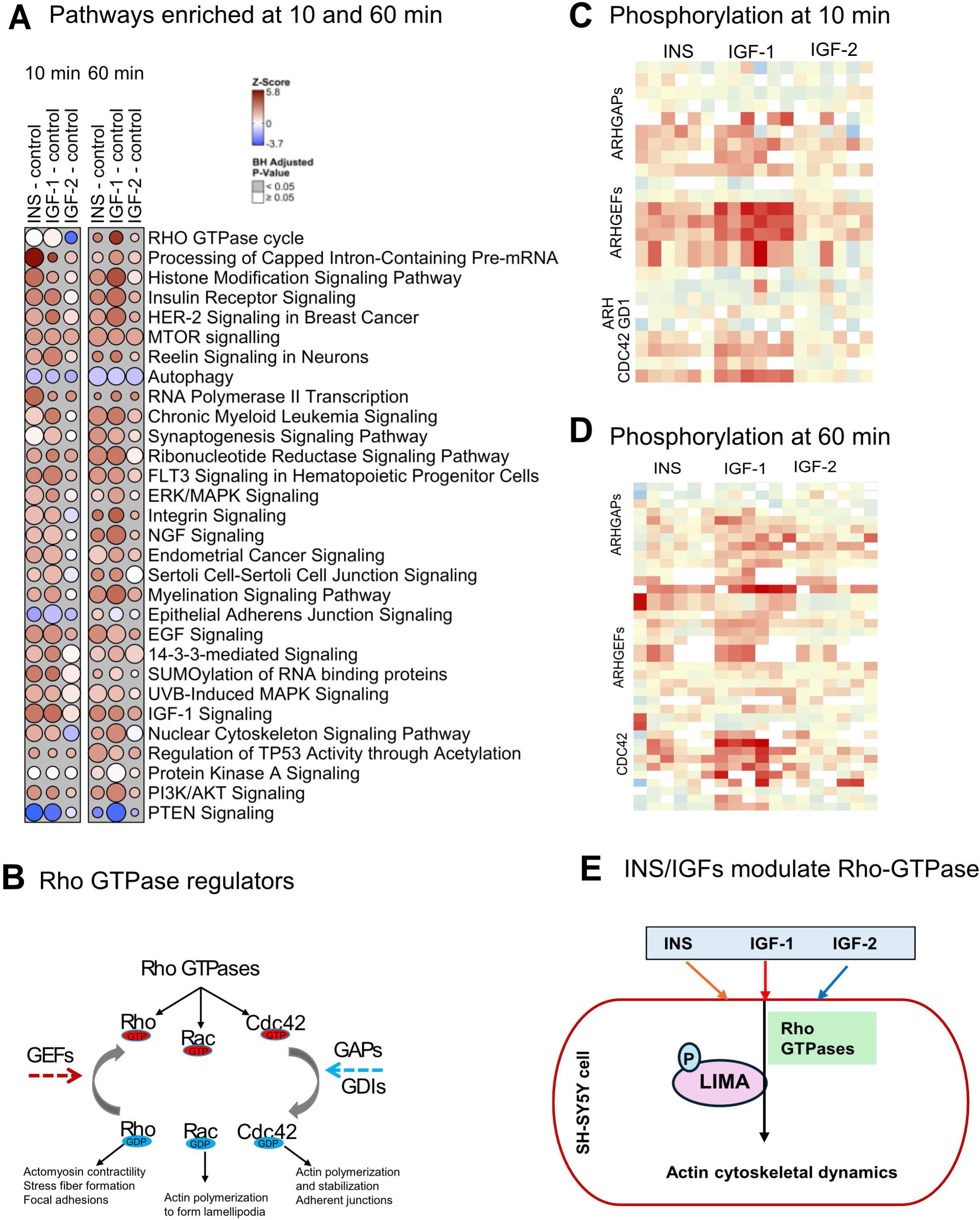
Pathway enrichment and analysis. (A) Bubble plot representing the top enriched pathways from IPA analysis of significant genes from INS, IGF-1 and IGF-2-stimulation at 10 min and 60 min compared to unstimulated controls. The color scale represents the z-score and the size of the bubble represents the fold-change. (B) Graphical summary of Rho GTPase regulators and effectors (C) Heatmap depicting Rho GTPase pathway components such as ARHGAPs, ARHGEFs, ARHGD1 & CDC42 and their phosphosites upon INS, IGF-1 and IGF-2 stimulation at 10 min compared to controls (D) Heatmap depicting Rho GTPase components and their phosphosites upon INS, IGF-1 and IGF-2 stimulation at 60min. (E) Summary of findings showing INS & IGFs may play a regulatory role in actin cytoskeletal dynamics through modulation of the Rho GTPase pathway.

### Rho GTPase Signaling dynamics

Rho GTPase signaling was identified as the most significant pathway across all ligands with an increase in pathway enrichment towards 60 min (Fig. 6A). Rho GTPase subunits Rho, RAC and CDC42 mediate the downstream activation of the pathway involving Guanine nucleotide exchange factor (GEFs) and Guanine nucleotide activator proteins and dissociation inhibitors (GAPs & GDIs) (Fig. 6B). We extracted the significant phosphosites of these subunits in our dataset (Fig. 6 C&D) and found that a more robust Rho signaling response is observed in IGF-1 stimulated cells compared to INS and IGF-2. ARHGAP-35, a GAP protein involved in neuronal morphogenesis^42^, is specifically phosphorylated by IGF-1. A similar increase was observed with RhoGEFs, ARHGEF12 and ARHGEF18, in IGF-1-stimulated cells, indicating that Rho GTPase components are predominantly phosphorylated by IGF-1. These results, together with the observed LIMA1-Y229 phosphorylation, suggest a role for phosphorylation events in regulating actin cytoskeletal dynamics by INS, IGF-1 and IGF-2 in neuronal cells (Figure 6E). We have provided the enrichment table listing the identified genes within each pathway from IPA analysis for further exploration (Supplementary dataset 2). This dataset enables the exploration of phosphosites from additional pathways, potentially linked to neuronal function, and supports hypothesis generation to broaden understanding of neuronal signaling in response to INS and IGF stimulation.

### IGF-1 regulates MRTFA expression via Rho GTPase pathway in SH-SY5Y cells

MRTFA is a known transcriptional target of Rho GTPase signaling^43^ and a coactivator of serum response factor (SRF), that plays a crucial role in cell growth and differentiation and modulates genes encoding cytoskeletal proteins^44^. Our data show an increase in phosphorylation of MRTFA-S6 upon INS, IGF-1 and IGF-2 stimulation at 10 min with a prominent response by IGF-1 (Fig. 7A). We did not observe a significant change in S6 phosphorylation at the 60 min time point (supplementary fig. 11). To test if IGF-1 mediates MRTFA expression via Rho GTPase signaling, we used chemical inhibitors such as Rhosin HCl (which inhibits the binding of RhoA to guanine nucleotide exchange factors, GEFs) and CCG 1423 (a Rho/SRF pathway inhibitor). We also included Wortmannin (a PI3K inhibitor) to determine whether MRTFA regulation operates directly under PI3K signaling. Western blot analysis of nuclear extract from cells stimulated with IGF-1 showed an increase in MRTFA expression (Fig. 7B & C), and this effect was abolished with CCG 1423 treatment and not by other inhibitors. Together, these results demonstrate that inhibiting the Rho/SRF pathway abolishes IGF-1-induced MRTFA expression in SH-SY5Y cells. Overall, our data suggests a role for INS and IGFs in phosphorylation modification of actin cytoskeletal protein dynamics and signaling in SH-SY5Y cells with IGF-1 showing a predominant response.

**Figure 7:**
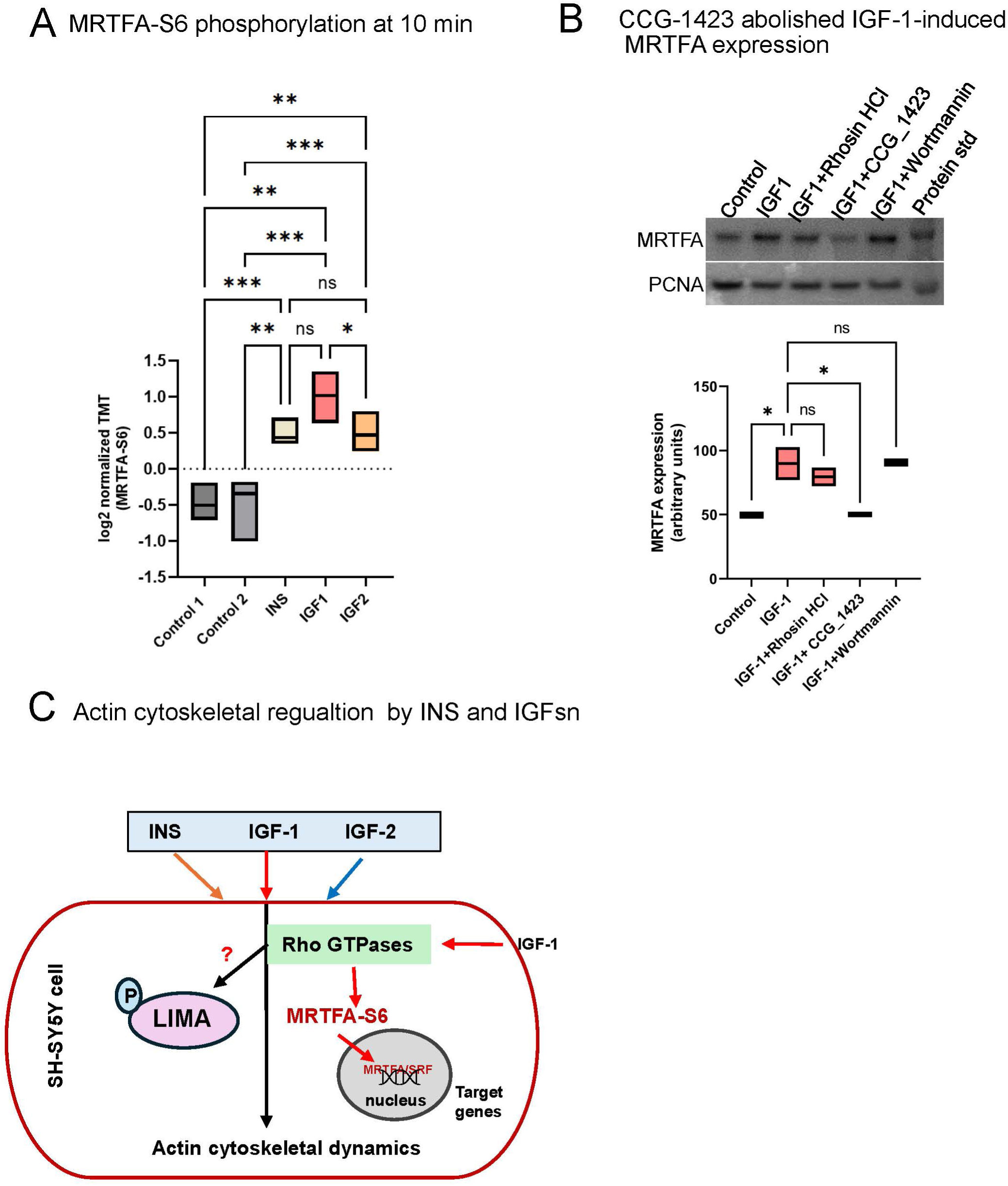
Rho GTPase and MRTFA regulation. (A) Abundance of MRTFA-S6 phosphosite identified from the phosphoproteomics data and represented as log2 transformed TMT value. One-way ANOVA was performed, * p < 0.05, ** p <0.005, *** p < 0.0005 and ns - not significant. (C) Validation of MRTFA regulation by IGF-1-mediate Rho GTPase was validated with western blotting. Nuclear extracts were collected from all groups and probed for MRTFA. Proliferative Cell nuclear Antigen (PCNA) was used as the nuclear internal control. Quantification of western blot bands was done using ImageJ software. We used N=2 independent experiments for the validation study. One-way ANOVA was used to determine the significance. * p < 0.05, ns – not significant. (D) Graphical summary if the role of INS and IGFs in regulating Rho GTPase and MRTFA expression

## Discussion

Our study revealed a comprehensive analysis of the phosphoproteome changes in SH-SY5Y cells in response to INS, IGF-1, and IGF-2 stimulation. The extensive dataset composed of 8,844 genes and 34,358 phosphosites, with 3,284 and 2,374 phosphosites showing significant changes at 10 and 60 minutes respectively, marks a pronounced dynamic shift over time. Notably, ligand-induced phosphorylation exhibited a decelerating trend at the 60-minute mark, reflecting the transient nature of these phosphorylation events. Despite a reduction in overall phosphosite numbers, IGF-1 showed an increase to 2,429 phosphosites compared to its count at 10 minutes. This suggests that INS and IGF-2 trigger immediate responses, whereas IGF-1 induces a more sustained phosphorylation profile. The individual and overlapped phosphosites among all ligands and across both time points underscores the complex, ligand-dependent temporal regulation of phosphorylation signaling. Phosphorylation of S, T, and Y residues displayed expected patterns, with S residues being the most prominent. Other studies have also showed low abundance of Y sites through phosphoproteomics studies^45–48^. A balanced level of Y phosphorylation is essential for maintaining cellular homeostasis, particularly in neuronal cells where dysregulation can contribute to neurodegenerative processes. Canonical receptors, INSR-Y1185, and IGF1R-Y1161, were identified and were significant at 10 minutes in INS and IGF-1 stimulated cells, with IGF-1 continuing their phosphorylation at 60 minutes. The presence of ALK-Y1507 indicates potential crosstalk with pathways related to cellular oncogenesis^49^. Since INS and IGF signaling are crucial for maintaining neuronal cell homeostasis, it is essential to identify the specific genes and phosphorylation sites stimulated by each ligand at 10 and 60 minutes. This understanding may shed light on the pathological mechanisms underlying dysregulated signaling, as INS and IGF-1 resistance^9^ and alteration in IGF-2 levels^50^ have been implicated in neurodegenerative diseases.

PATL-T194 was among the top phosphosite identified in all three ligands at both time points. The role of PATL in electrophysiology and transcriptional regulation of other genes has been studied in SH-SYSY neuroblastoma cell line and cardiomyocytes^51^. It directly regulates a human ether-a-go-go-related gene (hERG) K+ channel, which is essential for the normal electrical activity of cells^52^. Another set of phosphosites observed at 60 minutes exhibited similar trends across all ligands, including CAD-S1859 and STIP-S16. Ser1859 on CAD is a direct phosphorylation target of S6 kinase 1(S6K1), a downstream ribosomal protein target of mTORC1^53^. Phosphorylation at S1859 promotes CAD oligomerization, thereby enhancing de novo pyrimidine synthesis and facilitating S phase progression in mammalian cells. This cascade of phosphorylation is important for de novo pyrimidine synthesis and helps the overall progression of the cell cycle^54^. Our data support this by demonstrating cell cycle progression following stimulation with INS, IGF-1, and IGF-2. STIP1(Stress-induced phosphoprotein1)-S16 modification has been reported previously and this gene functions as a co-chaperone that plays a vital role in protein folding and cellular homeostasis, primarily by facilitating the interaction between the molecular chaperones Hsp70 and Hsp90^55^. STIP1 has been identified in a hypoglycemic model, underscoring its involvement in hypoglycemia-induced heat shock responses^56^. A Recent study reported that STIP1 plays a role in regulating the accumulation and toxicity of alpha synuclein in mouse models^57^. Therefore, the observed increase in STIP1-S16 phosphorylation following INS and IGF stimulation in neuronal cell lines warrants further investigation, as the S16 phosphorylation site may be critical to STIP1’s function in neurodegenerative diseases such as Lewy body disease (LBD) and Parkinson’s disease (PD), which are closely associated with alpha-synuclein pathology.

Apart from other canonical phosphorylation events, we also noticed an increase in phosphorylation of LIMA1-S225 at 10 min. Further exploration at the Y phosphorylation showed that LIMA1-Y229 is significantly high in all the ligands at 10 and 60 min. LIMA represents LIM domain and actin-binding protein 1 and is involved in actin cytoskeletal dynamics and thereby regulate cell shape and motility^58^. Identification of LIMA1 partners through STRING network provided insights into LIMA interactions with genes involved in actin cytoskeletal dynamics and Rho signaling^39,59^. FLNA and TJP both relate to cellular structures and processes by regulating cell shape and movement and tight junctional barriers respectively^60,61^. CTNNA1, B1 and D1 refers to Catenin proteins and are involved in cell adhesion and transcription regulation^62–64^. RhoA is a small GTPase associated with cytoskeletal organization. Given LIMA1’s interaction with these partners, it is evident that INS and IGFs regulate cell structure and cytoskeletal dynamics by phosphorylating LIMA1 and its associated targets. Maintaining structural integrity is vital for neuronal cells, as their highly polarized architecture underlies their ability to integrate, process, and transmit information. This is particularly relevant in the context of neurodegenerative diseases, where disruptions in cytoskeletal genes and Rho GTPase signaling have been implicated in the pathogenesis of disorders such as AD, PD and other conditions^65–67^. Given LIMA1’s critical role in membrane dynamics and metabolism^68^, it is important to further investigate the significance of its phosphorylation in neuronal cell function, particularly in response to INS and IGF stimulation.

FIP1L1-Y113 is another modified residue that showed a significant increase in INS and IGFs-treated cells at the 10-minute mark. It’s a pre-mRNA processing factor localized in the nucleus and functions together with cleavage and polyadenylation specificity factor (CPSF)^69^. As far as we know, phosphorylation at Y113 has not been previously reported and it is intriguing to further investigate the functional role of this phosphorylation modification in neuronal cells. We have also observed an increased phosphorylation of canonical MAPK signaling in INS and IGF-1 stimulated cells at 10 min. Given the association of exaggerated MAPK activity with neuronal apoptosis and the progression of AD pathology^70^, the role of INS and IGF-1 in regulating MAPK phosphorylation warrants closer investigation, especially since insulin and IGF-1 resistance has been found to be a key feature of AD^9,71–74^.

IPA analysis pointed us toward Rho GTPase signaling in this study. Rho GTPase function to regulate actin cytoskeletal dynamics^75^ and is quite significant in this context since aberrant Rho GTPases have been associated with cognitive dysfunction^76^.The detailed signaling network from this study identified many Guanine nucleotide exchange factors (GEFs) and GTPase-activating proteins (GAPs) including ARHGAP-35 which was notably high in IGF-1. ARHGAP is a Rho GTPase activating protein having known roles in cell migration, gene regulation and neuronal morphogenesis^42,77,78,79^. *ARHGAP35* is also a novel factor disrupted in human developmental eye phenotypes^42^. This reinforces IGF-1’s pivotal role in regulating actin cytoskeletal dynamics via Rho GTPase signaling and its role in maintaining the cellular plasticity. It also plays direct roles in neuronal morphogenesis and dendritic spine formation^80^. Additionally, ARHGAP35 is involved in the regulation of gene and mRNA translation through interactions with TFII-I and eIF3A^80^ and mice homozygous for a loss-of-function *Arhgap35* allele display structural brain anomalies^80^. Together, the role of Rho GTPase components in neuronal cell structure and function warrants further investigation, particularly in the context of INS and IGF stimulation.

Apart from the classical signaling pathways, we identified Reelin signaling in SH-SY5Y cells stimulated with all three ligands and a similar increased trend in IGF-1. Reelin is an extracellular, multifunctional signal glycoprotein that controls not only the positioning of neurons in the developing brain, but also their growth, maturation, and synaptic activity in the adult brain^81,82^. It is reported that Reelin signaling regulates actin-cytoskeletal dynamics by modulating Rho GTPases^83^. They bind to VLDLPR and APOER2 receptors and elicits downstream signaling^84^. Role of INS and IGFs in stimulating Reelin signaling, and their biological significance, functional targets and how thy crosstalk with Rho GTPase components in the brain remain unknown. APOER2 is a receptor protein interacts with APOE protein, encoded by the *APOE* gene and it is a strong genetic risk factor for AD^85^. Given Reelin’s interaction with APOER2, there is a pressing need to investigate how this signaling pathway is affected in INS and IGF-1 resistant forms of AD. Mapping the phosphorylation sites within this pathway could reveal new therapeutic targets.

One known target of Rho GTPase is MRTFA, a transcription coactivator that associates with the serum response factor (SRF) transcription factor involved in actin polymerization^43^. The SRF-MRTFA complex activity responds to Rho GTPase-induced changes in cellular globular actin (G-actin) concentration, thereby coupling cytoskeletal gene expression to cytoskeletal dynamics. Our data showed an increase in MRTFA phosphorylation at S6 upon stimulation with INS, IGF-1 and IGF-2. An increased response was observed with IGF-1, among the three ligands. We validated IGF-1-mediated MRTFA activation and nuclear localization in SH-SY5Y cells by a mechanism involving Rho GTPase pathway. These findings underscore the potential link between Rho GTPase signaling and MRTFA in mediating IGF-1’s effects on neuronal plasticity and survival. It would be valuable to investigate whether deficits in this pathway are present in INS/IGF-1-resistant neurodegenerative diseases. These findings highlight the therapeutic potential of modulating MRTFA in neurodegenerative diseases, where disrupted neuronal plasticity contributes to significant cognitive and motor impairments.

This study has a few limitations. First, the phosphoproteomics was performed on the SH-SY5Y cells. Although an established neuronal cell model, this is not a perfect representation of primary brain neurons, due to inherent genetic alterations and other limitations. Second, Although the dataset contains a wealth of information, we were unable to explore it in depth. Further network analysis is required to gain a comprehensive understanding of the phosphorylation dynamics specific to other significant pathways, including across cell types and time.

## Conclusions

The comprehensive phosphoproteomics analysis has provided the temporal and ligand-specific phosphorylation responses of INS, IGF-1 and IGF-2 in SH-SY5Y cells. The results have highlighted both the dynamics in phosphorylation events and the complex signaling pathways induced by these ligands. Specifically, phosphorylation of proteins associated with actin cytoskeletal dynamics, along with pathway enrichment analysis of Rho GTPase, revealed a link between the ligands and their role in actin remodeling in neuronal cells. Early stimulation of MRTFA-S6 phosphorylation by these ligands, coupled with the inhibition of MRTFA translocation by chemical inhibitors in IGF-1-stimulated cells, provided further evidence supporting this relationship. The study also generated several testable hypotheses which can be further explored to better understand the roles of INS and IGFs in neuronal physiology and pathology, including neurodegenerative processes.

## List of abbreviations

INS: Insulin
IGF-1: Insulin-like growth factor-1
IGF-2: Insulin-like growth factor-2
IIS: Insulin/IGF signaling
IR: Insulin receptor
IGF-1R: Insulin-like growth factor 1 receptor
ALK: Anaplastic lymphoma kinase
IMAC: Immobilized metal affinity chromatography
TMT: Tandem mass tag
SPE: Solid phase extraction
SDC: sodium deoxycholate
TCEP: tris(2-carboxyethyl) phosphine
CAM: 2-chloroacetamide
TFA: Trifluoroacetic acid
FA: Formic acid
PCNA: Proliferating cell nuclear antigen
LC-MS/MS: Liquid chromatography-tandem mass spectrometry
IPA: Ingenuity pathway analysis
LIMA1: LIM domain and actin-binding protein 1
SRF: Serum response factor
STIP1: Stress-induced phosphoprotein1
CPSF: Cleavage and polyadenylation specificity factor
Rho: GTPase Ras homology guanosine triphosphate
MRTFA: Myocardin related transcription factor A
ARHGEFs: Rho Guanine nucleotide exchange factors
ARHGAPs: Rho GTPase activating proteins
PI3K: Phosphoinositide 3-kinase
MAPK: Mitogen-activated protein kinase
AD: Alzheimer’s disease
PD: Parkinson’s disease
LBD: Lewy body disease

## Declarations

### Ethics approval and consent to participate

Not applicable

### Consent for publication

Not applicable

### Availability of data and materials

All raw files have been deposited in the MassIVE data repository and are accessible with the following ID: MSV000097782. All data generated or analysed during the current study are included in the supplementary files.

### Competing interests

The authors declare that they have no competing interests

### Funding

This work was supported by the NIH National Institute of Aging grants U01 AG061356 (to P.L.D.J) and R01 AG074549 (to Z.A). Part of this work was performed in the Environmental Molecular Science Laboratory, a U.S. Department of Energy (DOE) national scientific user facility at Pacific Northwest National Laboratory (PNNL). Battelle operates PNNL for the DOE under contract DE-AC05-76RLO01830. The opinions expressed in this article are the authors’ own and do not reflect the view of the NIH, the Department of Health and Human Services, or the U.S. government.

### Authors’ contributions

MGU performed data interpretation and analysis, conducted validation experiments, wrote and reviewed the manuscript. RS performed preliminary cell culture experiments and phosphoproteomics sample preparation. CP conducted data analysis. MT Contributed to the methodology and provided experimental support. HO provided experimental support. PDJ **r**eviewed the manuscript. DB reviewed the manuscript. ZA provided constructive feedback and reviewed the manuscript. VAP contributed to project conceptualization, performed data analysis, provided supervision, edited and reviewed the manuscript.

## Acknowledgements

The authors thank Tyler J. Sagendorf for writing the code to generate bubble heatmaps and Matthew Monroe for assistance with depositing the raw phosphoproteomics data to the MassIVE repository.

## Notes

### Competing Interest Statement

The authors have declared no competing interest.

## References

1. Cai, W. et al. Domain-dependent effects of insulin and IGF-1 receptors on signalling and gene expression. Nat Commun 8, 14892 (2017).

2. Okawa, E. R. et al. Essential roles of insulin and IGF-1 receptors during embryonic lineage development. Mol Metab 47, 101164 (2021).

3. Nagao, H. et al. Distinct signaling by insulin and IGF-1 receptors and their extra- and intracellular domains. Proceedings of the National Academy of Sciences 118, e2019474118 (2021).

4. LeRoith, D., Holly, J. M. P. & Forbes, B. E. Insulin-like growth factors: Ligands, binding proteins, and receptors. Molecular Metabolism 52, 101245 (2021).

5. O’Neill, B. T. et al. Differential Role of Insulin/IGF-1 Receptor Signaling in Muscle Growth and Glucose Homeostasis. Cell Reports 11, 1220–1235 (2015).

6. Massoner, P., Ladurner-Rennau, M., Eder, I. E. & Klocker, H. Insulin-like growth factors and insulin control a multifunctional signalling network of significant importance in cancer. Br J Cancer 103, 1479–1484 (2010).

7. Milstein, J. L. & Ferris, H. A. The brain as an insulin-sensitive metabolic organ. Mol Metab 52, 101234 (2021).

8. Blázquez, E., Velázquez, E., Hurtado-Carneiro, V. & Ruiz-Albusac, J. M. Insulin in the Brain: Its Pathophysiological Implications for States Related with Central Insulin Resistance, Type 2 Diabetes and Alzheimer’s Disease. Front Endocrinol (Lausanne) 5, 161 (2014).

9. Talbot, K. et al. Demonstrated brain insulin resistance in Alzheimer’s disease patients is associated with IGF-1 resistance, IRS-1 dysregulation, and cognitive decline. J Clin Invest 122, 1316–1338 (2012).

10. Rhea, E. M. et al. State of the Science on Brain Insulin Resistance and Cognitive Decline Due to Alzheimer’s Disease. Aging Dis 15, 1688–1725 (2024).

11. Chen, W., Cai, W., Hoover, B. & Kahn, C. R. Insulin action in the brain: cell types, circuits, and diseases. Trends in Neurosciences 45, 384–400 (2022).

12. Dyer, A. H., Vahdatpour, C., Sanfeliu, A. & Tropea, D. The role of Insulin-Like Growth Factor 1 (IGF- 1) in brain development, maturation and neuroplasticity. Neuroscience 325, 89–99 (2016).

13. Alberini, C. M. IGF2 in memory, neurodevelopmental disorders, and neurodegenerative diseases. Trends Neurosci 46, 488–502 (2023).

14. Qatanani, M. & Lazar, M. A. Mechanisms of obesity-associated insulin resistance: many choices on the menu. Genes Dev. 21, 1443–1455 (2007).

15. Gan, L. et al. Static training improves insulin resistance in skeletal muscle of type 2 diabetic mice via the IGF-2/IGF-1R pathway. Sci Rep 15, 10662 (2025).

16. Li, M. et al. Trends in insulin resistance: insights into mechanisms and therapeutic strategy. Sig Transduct Target Ther 7, 1–25 (2022).

17. Brown, A. E. & Walker, M. Genetics of Insulin Resistance and the Metabolic Syndrome. Curr Cardiol Rep 18, 75 (2016).

18. Maratou, E. et al. Studies of insulin resistance in patients with clinical and subclinical hypothyroidism. European Journal of Endocrinology 160, 785–790 (2009).

19. Manrique, C., Johnson, M. & Sowers, J. R. Thiazide Diuretics Alone or with Beta-blockers Impair Glucose Metabolism in Hypertensive Patients with Abdominal Obesity. Hypertension 55, 15–17 (2010).

20. Geer, E. B., Islam, J. & Buettner, C. Mechanisms of Glucocorticoid-Induced Insulin Resistance. Endocrinol Metab Clin North Am 43, 75–102 (2014).

21. Kolb, H., Kempf, K. & Martin, S. Insulin and aging – a disappointing relationship. Front Endocrinol (Lausanne) 14, 1261298 (2023).

22. Arvanitakis, Z., Wilson, R. S., Bienias, J. L., Evans, D. A. & Bennett, D. A. Diabetes mellitus and risk of Alzheimer disease and decline in cognitive function. Arch Neurol 61, 661–666 (2004).

23. Arvanitakis, Z. et al. Brain Insulin Signaling, Alzheimer Disease Pathology, and Cognitive Function. Ann Neurol 88, 513–525 (2020).

24. Zhao, W.-Q. & Townsend, M. Insulin resistance and amyloidogenesis as common molecular foundation for type 2 diabetes and Alzheimer’s disease. Biochimica et Biophysica Acta (BBA) - Molecular Basis of Disease 1792, 482–496 (2009).

25. Desch, K., Schuman, E. M. & Langer, J. D. Quantifying phosphorylation dynamics in primary neuronal cultures using LC-MS/MS. STAR Protocols 3, 101063 (2022).

26. Jiang, W. et al. Phosphoproteome Analysis Identifies a Synaptotagmin-1-Associated Complex Involved in Ischemic Neuron Injury. Mol Cell Proteomics 21, 100222 (2022).

27. Nakamura, M. et al. Phosphoproteomic profiling of human SH-SY5Y neuroblastoma cells during response to 6-hydroxydopamine-induced oxidative stress. Biochim Biophys Acta 1763, 977–989 (2006).

28. Leung, T. C. N. et al. Temporal Quantitative Proteomic and Phosphoproteomic Profiling of SH- SY5Y and IMR-32 Neuroblastoma Cells during All-Trans-Retinoic Acid-Induced Neuronal Differentiation. Int J Mol Sci 25, 1047 (2024).

29. Biedler, J. L., Helson, L. & Spengler, B. A. Morphology and Growth, Tumorigenicity, and Cytogenetics of Human Neuroblastoma Cells in Continuous Culture1. Cancer Research 33, 2643–2652 (1973).

30. Image Analysis and Quantitation for Western Blotting | Bio-Rad. https://www.bio-rad.com/en-us/applications-technologies/image-analysis-quantitation-for-western-blotting?ID=PQEERM9V5F6X.

31. Humphrey, S. J., Azimifar, S. B. & Mann, M. High-throughput phosphoproteomics reveals in vivo insulin signaling dynamics. Nat Biotechnol 33, 990–995 (2015).

32. Humphrey, S. J., Karayel, O., James, D. E. & Mann, M. High-throughput and high-sensitivity phosphoproteomics with the EasyPhos platform. Nat Protoc 13, 1897–1916 (2018).

33. Mertins, P. et al. Reproducible workflow for multiplexed deep-scale proteome and phosphoproteome analysis of tumor tissues by liquid chromatography–mass spectrometry. Nat Protoc 13, 1632–1661 (2018).

34. Kim, S. & Pevzner, P. A. MS-GF+ makes progress towards a universal database search tool for proteomics. Nat Commun 5, 5277 (2014).

35. Monroe, M. E., Shaw, J. L., Daly, D. S., Adkins, J. N. & Smith, R. D. MASIC: a software program for fast quantitation and flexible visualization of chromatographic profiles from detected LC-MS(/MS) features. Comput Biol Chem 32, 215–217 (2008).

36. The UniProt Consortium. UniProt: the Universal Protein Knowledgebase in 2025. Nucleic Acids Research 53, D609–D617 (2025).

37. Johnson, W. E., Li, C. & Rabinovic, A. Adjusting batch effects in microarray expression data using empirical Bayes methods. Biostatistics 8, 118–127 (2007).

38. Sagendorf, T. TMSig: Tools for Molecular Signatures. https://bioconductor.org/packages/TMSig(2024) doi:doi:10.18129/B9.bioc.TMSig.

39. Szklarczyk, D. et al. The STRING database in 2023: protein-protein association networks and functional enrichment analyses for any sequenced genome of interest. Nucleic Acids Res 51, D638–D646 (2023).

40. Hu, J., et al. Opposing FlnA and FlnB interactions regulate RhoA activation in guiding dynamic actin stress fiber formation and cell spreading. Hum Mol Genet 26, 1294–1304 (2017).

41. Escobar, D. J., et al. α-Catenin phosphorylation promotes intercellular adhesion through a dual-kinase mechanism. J Cell Sci 128, 1150–1165 (2015).

42. Reis, L. M., et al. ARHGAP35 is a novel factor disrupted in human developmental eye phenotypes. Eur J Hum Genet 31, 363–367 (2023).

43. Esnault, C., et al. Rho-actin signaling to the MRTF coactivators dominates the immediate transcriptional response to serum in fibroblasts. Genes Dev 28, 943–958 (2014).

44. Reed, F., Larsuel, S. T., Mayday, M. Y., Scanlon, V. & Krause, D. S. MRTFA: A critical protein in normal and malignant hematopoiesis and beyond. Journal of Biological Chemistry 296, 100543 (2021).

45. Hunter, T. & Sefton, B. M. Transforming gene product of Rous sarcoma virus phosphorylates tyrosine. Proceedings of the National Academy of Sciences 77, 1311–1315 (1980).

46. Gerritsen, J. S. & White, F. M. Phosphoproteomics: a valuable tool for uncovering molecular signaling in cancer cells. Expert Rev Proteomics 18, 661–674 (2021).

47. Heibeck, T. H., et al. An Extensive Survey of Tyrosine Phosphorylation Revealing New Sites in Human Mammary Epithelial Cells. J Proteome Res 8, 3852–3861a (2009).

48. Wolf-Yadlin, A., Hautaniemi, S., Lauffenburger, D. A. & White, F. M. Multiple reaction monitoring for robust quantitative proteomic analysis of cellular signaling networks. Proc Natl Acad Sci U S A 104, 5860–5865 (2007).

49. Huang, H., et al. Extracellular domain shedding of the ALK receptor mediates neuroblastoma cell migration. Cell Rep 36, 109363 (2021).

50. Hertze, J., Nägga, K., Minthon, L. & Hansson, O. Changes in cerebrospinal fluid and blood plasma levels of IGF-II and its binding proteins in Alzheimer’s disease: an observational study. BMC Neurol 14, 64 (2014).

51. Yao, L., Ruan, M.-Y., Ye, S.-W. & Cai, S.-Q. DNA topoisomerase 2-associated proteins PATL1 and PATL2 regulate the biogenesis of hERG K+ channels. Proc Natl Acad Sci U S A 120, e2206146120 (2023).

52. Cui, Z. & Zhang, S. Regulation of the Human Ether-a-go-go-related Gene (hERG) Channel by Rab4 Protein through Neural Precursor Cell-expressed Developmentally Down-regulated Protein 4-2 (Nedd4-2). J Biol Chem 288, 21876–21886 (2013).

53. Ben-Sahra, I., Howell, J. J., Asara, J. M. & Manning, B. D. Stimulation of de novo pyrimidine synthesis by growth signaling through mTOR and S6K1. Science 339, 1323–1328 (2013).

54. Robitaille, A. M., et al. Quantitative phosphoproteomics reveal mTORC1 activates de novo pyrimidine synthesis. Science 339, 1320–1323 (2013).

55. da Fonseca, A. C. C., et al. The multiple functions of the co-chaperone stress inducible protein 1. Cytokine & Growth Factor Reviews 57, 73–84 (2021).

56. Atkin, A. S., et al. Impact of severe hypoglycemia on the heat shock and related protein response. Sci Rep 11, 17057 (2021).

57. Lackie, R. E., et al. Stress-inducible phosphoprotein 1 (HOP/STI1/STIP1) regulates the accumulation and toxicity of α-synuclein in vivo. Acta Neuropathol 144, 881–910 (2022).

58. Wang, X., et al. Characterization of LIMA1 and its emerging roles and potential therapeutic prospects in cancers. Front Oncol 13, 1115943 (2023).

59. Szklarczyk, D. et al. The STRING database in 2021: customizable protein-protein networks, and functional characterization of user-uploaded gene/measurement sets. Nucleic Acids Res 49, D605–D612 (2021).

60. Noam, Y., et al. Filamin A Promotes Dynamin-dependent Internalization of Hyperpolarization-activated Cyclic Nucleotide-gated Type 1 (HCN1) Channels and Restricts Ih in Hippocampal Neurons. J Biol Chem 289, 5889–5903 (2014).

61. González-Mariscal, L., Betanzos, A., Nava, P. & Jaramillo, B. E. Tight junction proteins. Prog Biophys Mol Biol 81, 1–44 (2003).

62. van der Wal, T. & van Amerongen, R. Walking the tight wire between cell adhesion and WNT signalling: a balancing act for β-catenin. Open Biol 10, 200267 (2020).

63. Wang, J., et al. A Ctnnb1 enhancer regulates neocortical neurogenesis by controlling the abundance of intermediate progenitors. Cell Discov 8, 1–17 (2022).

64. Herrera-Pariente, C., et al. CTNND1 is involved in germline predisposition to early-onset gastric cancer by affecting cell-to-cell interactions. Gastric Cancer 27, 747–759 (2024).

65. Mendoza-Naranjo, A., Gonzalez-Billault, C. & Maccioni, R. B. Aβ1-42 stimulates actin polymerization in hippocampal neurons through Rac1 and Cdc42 Rho GTPases. Journal of Cell Science 120, 279–288 (2007).

66. Zhou, Z., et al. Rho GTPase regulation of α-synuclein and VMAT2: implications for pathogenesis of Parkinson’s disease. Mol Cell Neurosci 48, 29–37 (2011).

67. Tourette, C., et al. A large scale Huntingtin protein interaction network implicates Rho GTPase signaling pathways in Huntington disease. J Biol Chem 289, 6709–6726 (2014).

68. Duethorn, B., et al. Lima1 mediates the pluripotency control of membrane dynamics and cellular metabolism. Nat Commun 13, 610 (2022).

69. Kaufmann, I., Martin, G., Friedlein, A., Langen, H. & Keller, W. Human Fip1 is a subunit of CPSF that binds to U-rich RNA elements and stimulates poly(A) polymerase. EMBO J 23, 616–626 (2004).

70. Zhu, X., Lee, H., Raina, A. K., Perry, G. & Smith, M. A. The role of mitogen-activated protein kinase pathways in Alzheimer’s disease. Neurosignals 11, 270–281 (2002).

71. Arnold, S. E., et al. Brain insulin resistance in type 2 diabetes and Alzheimer disease: concepts and conundrums. Nat Rev Neurol 14, 168–181 (2018).

72. Tong, H., et al. Brain Insulin Signaling is Associated with Late-Life Cognitive Decline. Aging and disease 15, 2205–2215 (2024).

73. Tong, H., et al. Alzheimer’s disease-related cortical proteins modify the association of brain insulin signaling with cognitive decline. Journal of Alzheimer’s Disease 104, 667–677 (2025).

74. Moloney, A. M., et al. Defects in IGF-1 receptor, insulin receptor and IRS-1/2 in Alzheimer’s disease indicate possible resistance to IGF-1 and insulin signalling. Neurobiology of Aging 31, 224–243 (2010).

75. Hall, A. Rho GTPases and the actin cytoskeleton. Science 279, 509–514 (1998).

76. De Filippis, B., Romano, E. & Laviola, G. Aberrant Rho GTPases signaling and cognitive dysfunction: In vivo evidence for a compelling molecular relationship. Neuroscience & Biobehavioral Reviews 46, 285–301 (2014).

77. Govek, E.-E., Hatten, M. E. & Van Aelst, L. The Role of Rho GTPase Proteins in CNS Neuronal Migration. Dev Neurobiol 71, 528–553 (2011).

78. Héraud, C., Pinault, M., Lagrée, V. & Moreau, V. p190RhoGAPs, the ARHGAP35- and ARHGAP5- Encoded Proteins, in Health and Disease. Cells 8, 351 (2019).

79. Trompeter, N., et al. Insulin-like Growth Factor-1 Regulates the Mechanosensitivity of Chondrocytes by Modulating TRPV4. Cell Calcium 99, 102467 (2021).

80. Brouns, M. R., et al. The adhesion signaling molecule p190 RhoGAP is required for morphogenetic processes in neural development. Development 127, 4891–4903 (2000).

81. Stranahan, A. M., Haberman, R. P. & Gallagher, M. Cognitive decline is associated with reduced reelin expression in the entorhinal cortex of aged rats. Cereb Cortex 21, 392–400 (2011).

82. Stranahan, A. M., Erion, J. R. & Wosiski-Kuhn, M. Reelin signaling in development, maintenance, and plasticity of neural networks. Ageing Res Rev 12, 815–822 (2013).

83. Leemhuis, J. & Bock, H. H. Reelin modulates cytoskeletal organization by regulating Rho GTPases. Commun Integr Biol 4, 254–257 (2011).

84. Andersen, O. M., Benhayon, D., Curran, T. & Willnow, T. E. Differential binding of ligands to the apolipoprotein E receptor 2. Biochemistry 42, 9355–9364 (2003).

85. Serrano-Pozo, A., Das, S. & Hyman, B. T. APOE and Alzheimer’s Disease: Advances in Genetics, Pathophysiology, and Therapeutic Approaches. Lancet Neurol 20, 68–80 (2021).

