## Supplementary figures and images for "Phosphoproteomics unveils the signaling dynamics in neuronal cells stimulated with insulin and insulin-like growth factors"

### Supplementary figure 2

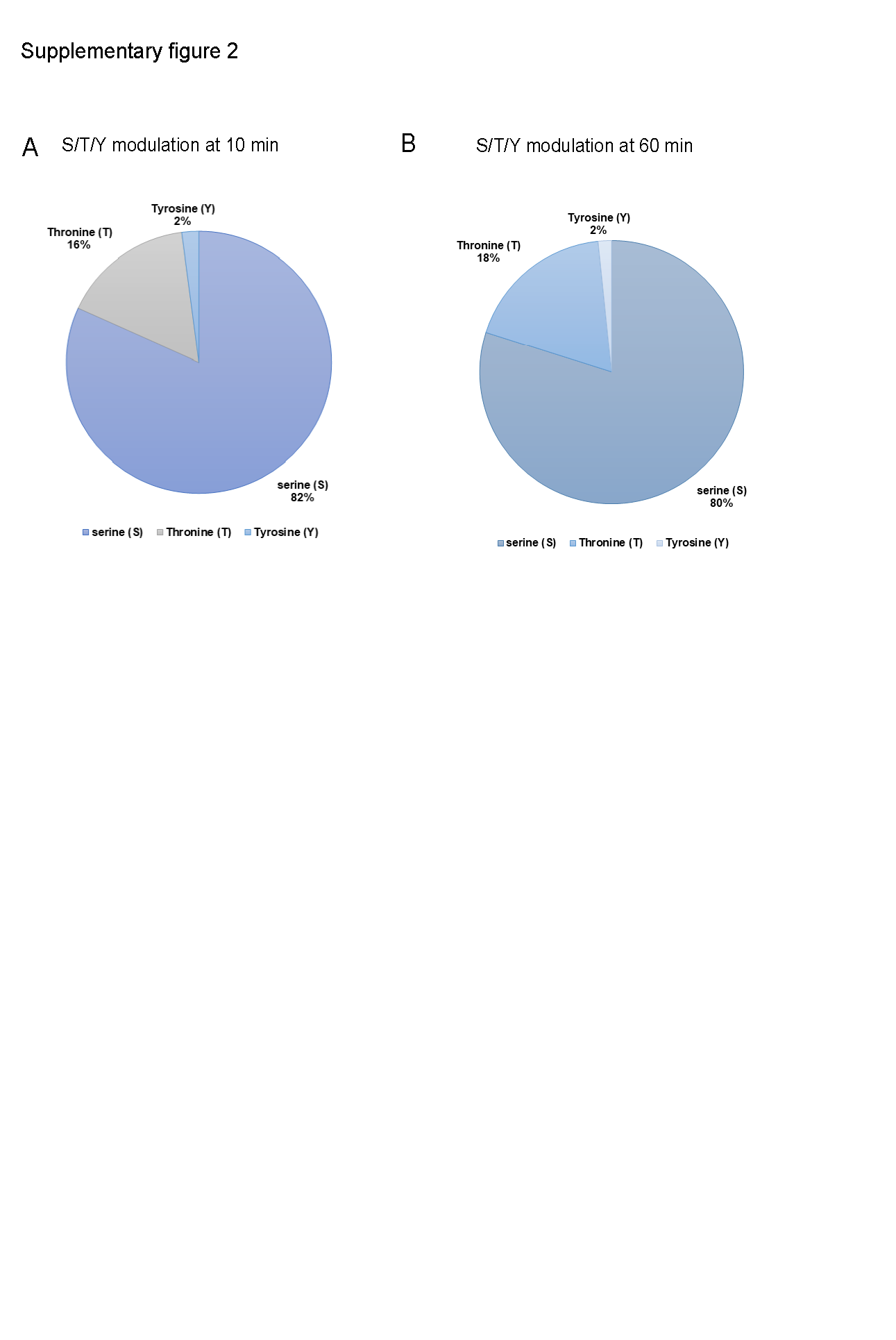

### Supplementary figure 3

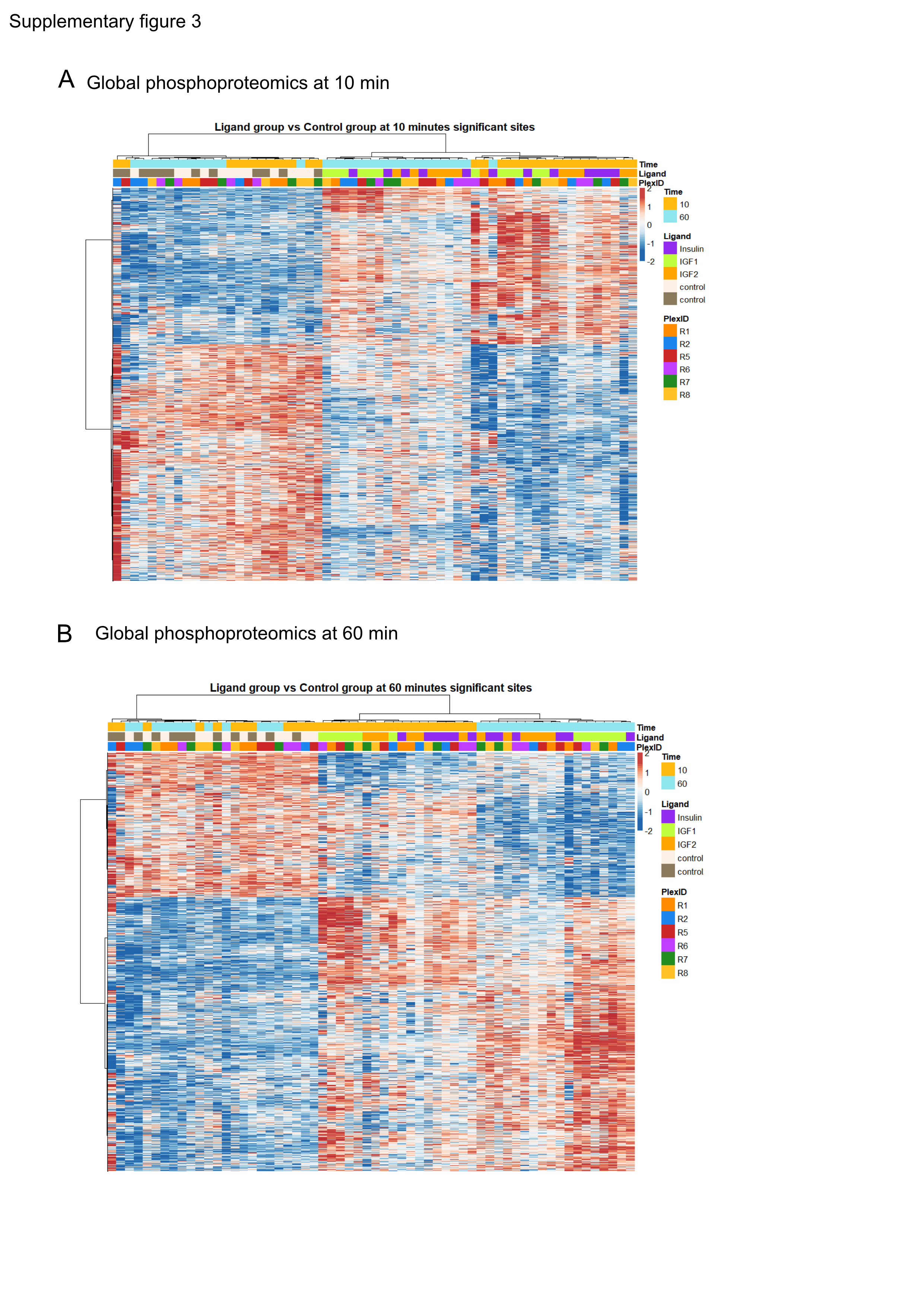

### Supplementary figure 4

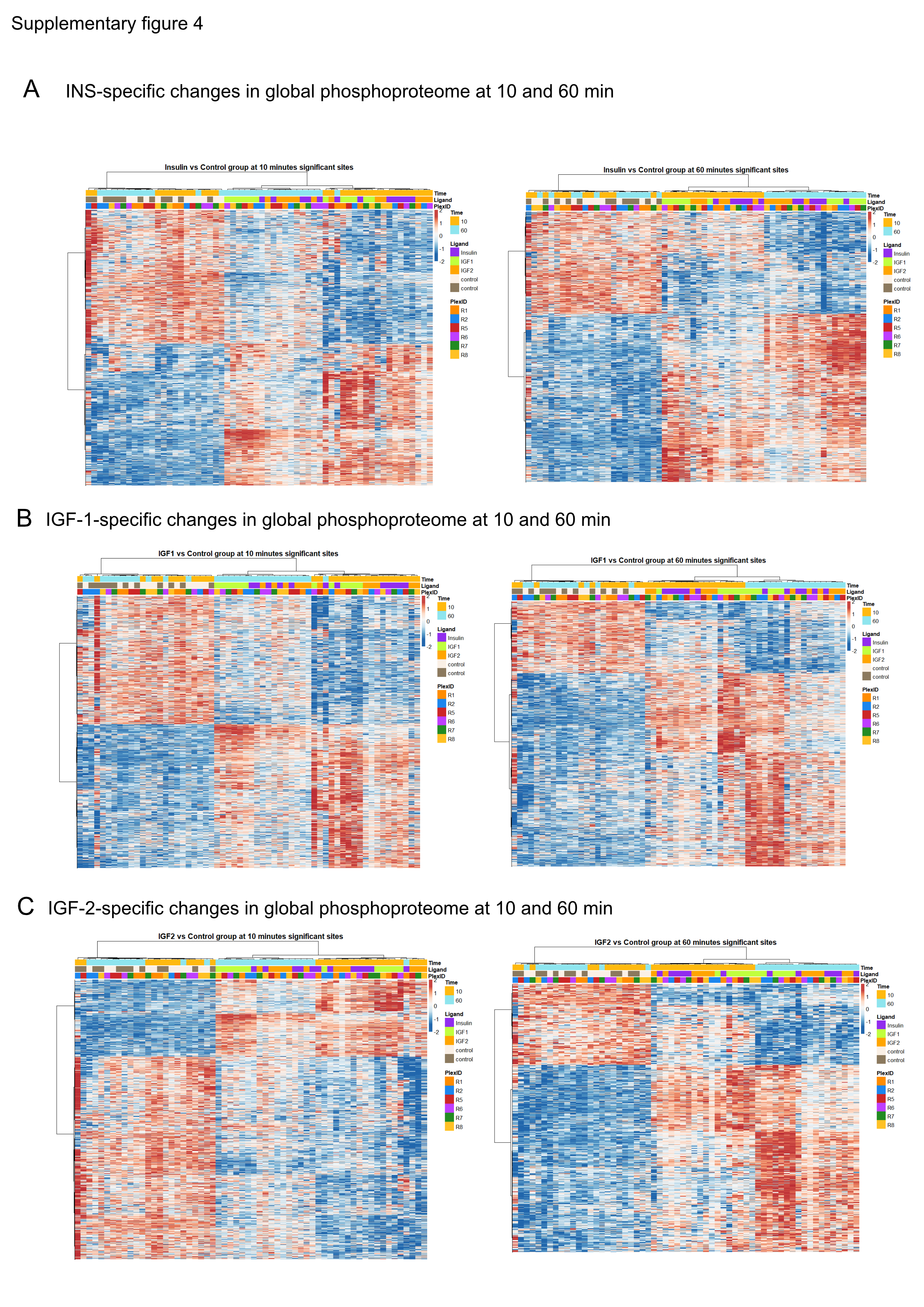

### Supplementary figure 5

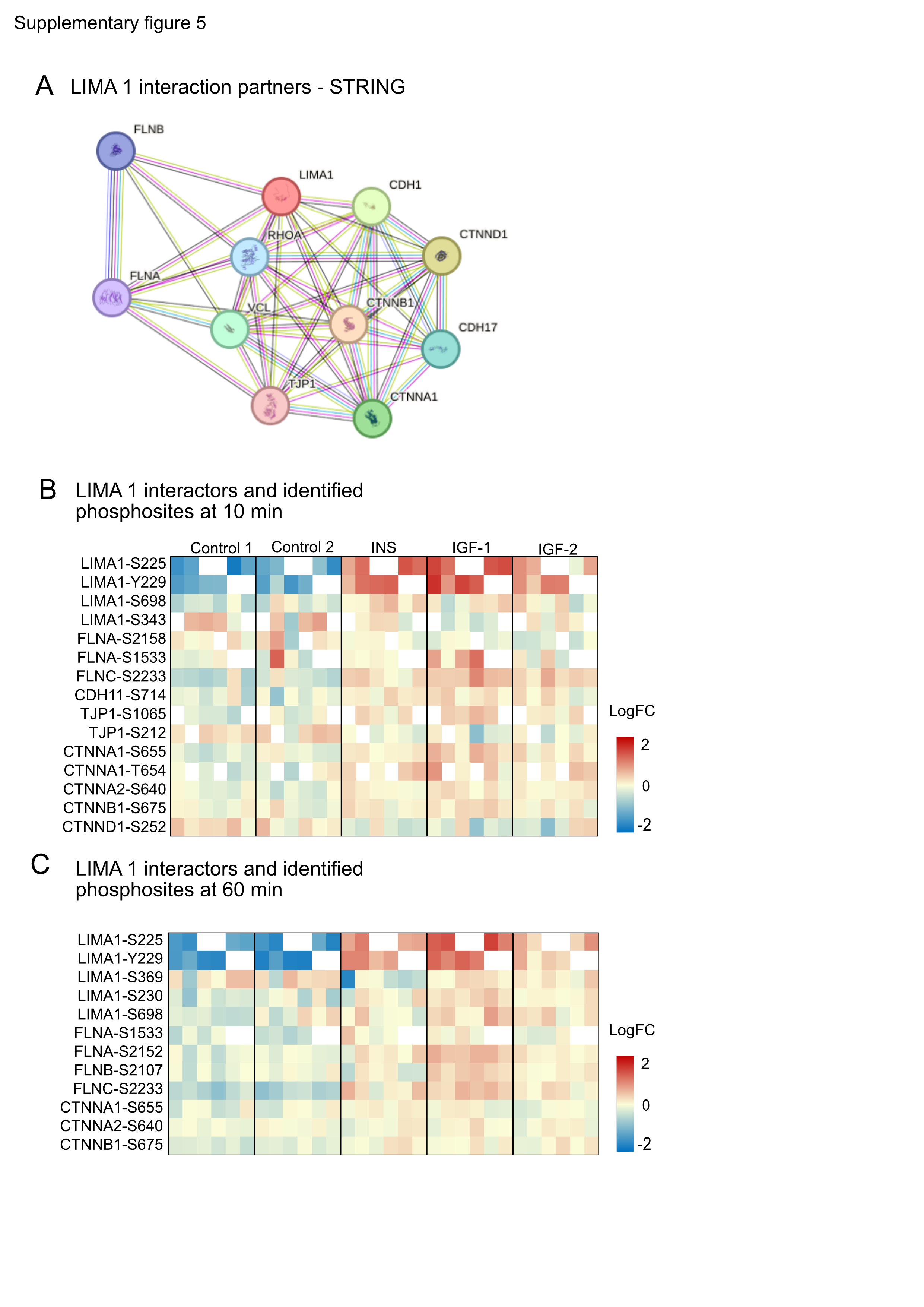

### Supplementary figure 6

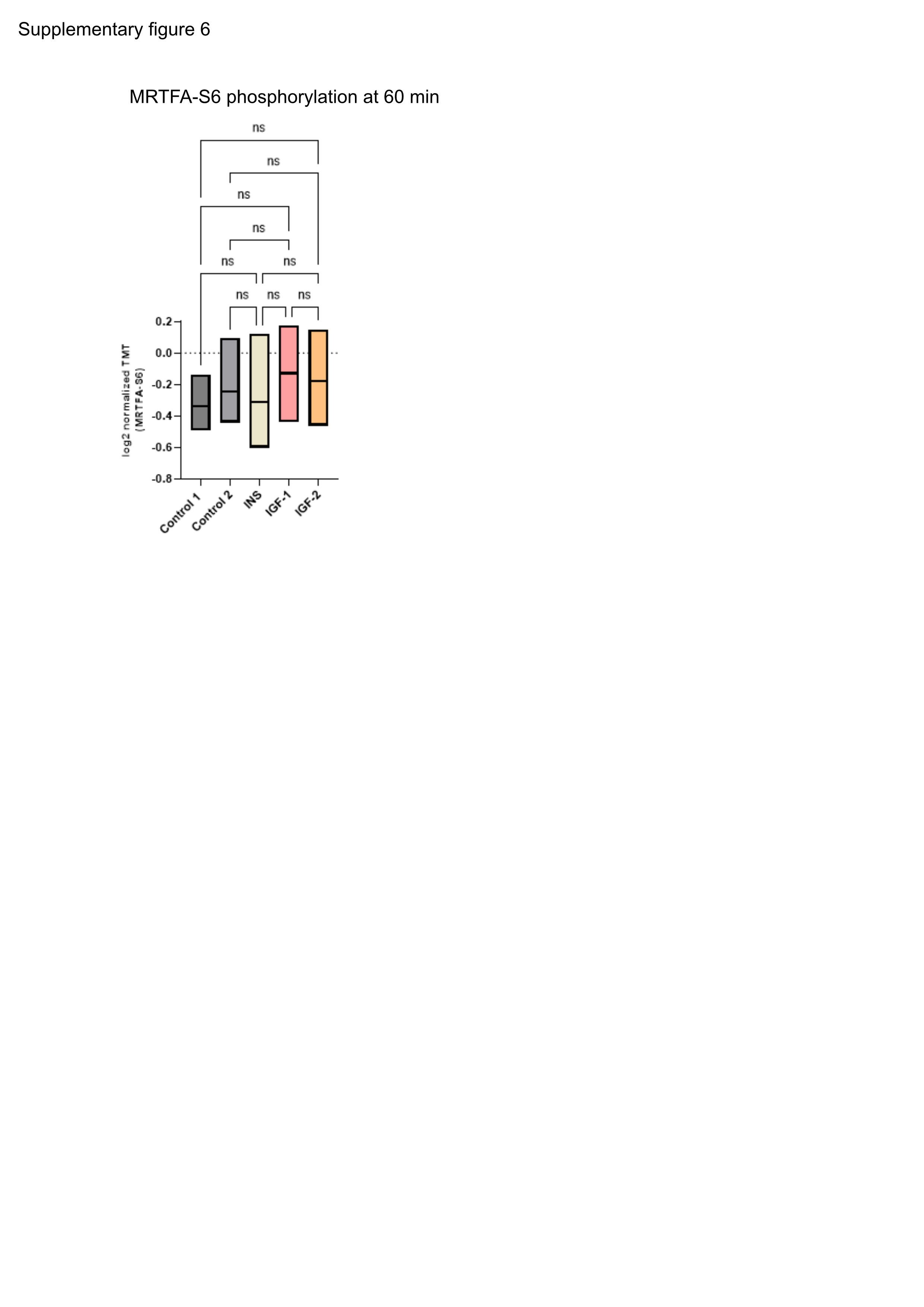

### Supplentary figure 1

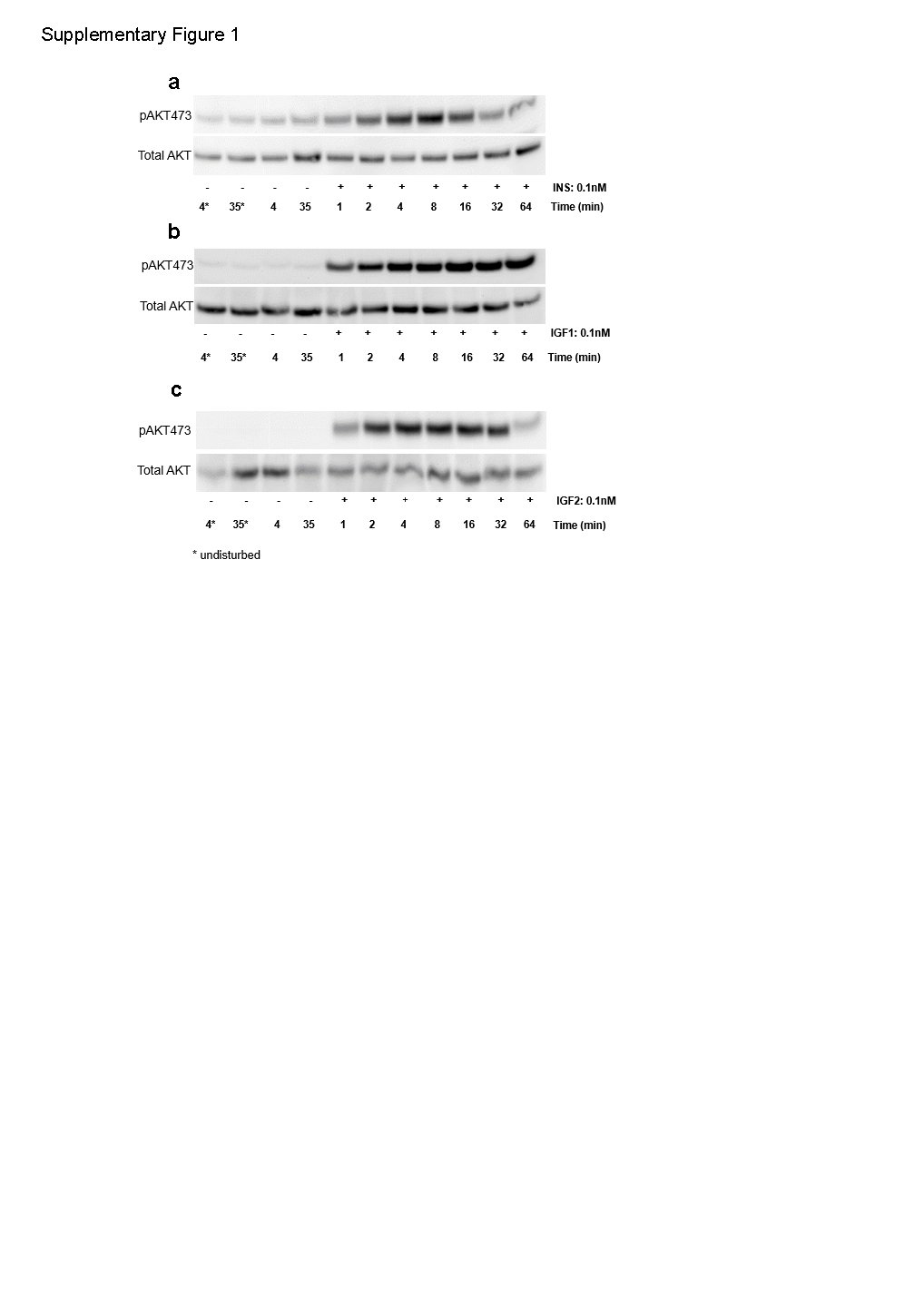
